# Zika virus impacts extracellular vesicle composition and cellular gene expression in macaque early gestation trophoblasts

**DOI:** 10.1101/2021.10.07.463494

**Authors:** Lindsey N. Block, Jenna Kropp Schmidt, Megan C. McKeon, Brittany D. Bowman, Gregory J. Wiepz, Thaddeus G. Golos

## Abstract

Zika virus (ZIKV) infection at the maternal-placental interface is associated with adverse pregnancy outcomes including fetal demise and pregnancy loss. To determine how infection impacts placental trophoblasts, we utilized rhesus macaque trophoblast stem cells (TSC) that can be differentiated into early gestation syncytiotrophoblasts (ST) and extravillous trophoblasts (EVT). TSCs and STs, but not EVTs, were highly permissive to productive infection with ZIKV strain DAK AR 41524. The impact of ZIKV on the cellular transcriptome showed that infection of TSCs and STs increased expression of immune related genes, including those involved in type I and type III interferon responses. ZIKV exposure altered extracellular vesicle (EV) protein, mRNA, and miRNA cargo, regardless of productive infection. These findings suggest that early gestation macaque TSCs and STs are permissive to ZIKV infection, and that EV analysis may provide a foundation for identifying non-invasive biomarkers of placental infection in a highly translational model.

## Introduction

Maternal Zika virus (ZIKV) infection is associated with adverse pregnancy outcomes including pregnancy loss and fetal malformations ^1,2^. Prolonged maternal viremia in humans and nonhuman primates ^2-7^ suggests there is a pregnancy-specific viral reservoir, and the presence of virus at the maternal fetal interface ^8^ support the premise that the placenta may serve as a viral reservoir. Thus, there is a pressing need to better understand the mechanisms of placental infection and ZIKV impact on placental cell function.

The placenta is an essential organ in pregnancy as it not only conveys sufficient nutrients and oxygen for proper development ^9^, but also provides signals for the adaptation of maternal physiological systems to pregnancy. Errors in placental development and function are often associated with adverse pregnancy outcomes ^9^. The placenta is comprised of several trophoblast cell types, including villous cytotrophoblasts (vCTBs), syncytiotrophoblast (STs), extravillous trophoblasts (EVTs), as well as fetal macrophages (Hofbauer cells), other immune and vascular cells, and fibroblasts ^10^. vCTBs can be programmed to a proliferative stem cell-like state in vitro, termed trophoblast stem cells (TSCs) that can be differentiated into STs or EVTs ^10-12^.

Maternal ZIKV infection early in pregnancy is associated with increased risk for pregnancy loss and more severe complications ^2,4,13-17^. However, the ability of clinicians and investigators to assess the risk of vertical transmission is hindered by uncertainty in the timing of infection and the inability to directly assess placental, fetal, or reproductive tract tissues during pregnancy. In addition, in vitro studies have demonstrated that early gestation trophoblasts are permissive to ZIKV infection; however, the specific impact of infection on trophoblasts has not been comprehensively characterized. Inconsistencies in in vivo reports of trophoblast infection warrants further study of trophoblast-type permissiveness to ZIKV infection in early pregnancy to enable future development of methods to non-invasively monitor in vivo placental infection. Extracellular vesicles (EVs) are widely studied as a minimally invasive “liquid biopsy” to monitor cell function from in vivo body fluids or in vitro culture media ^18,19^. EVs contain cell-type specific markers, which can be used to monitor the status of specific cells ^18^. The cargo packaged within EVs, including nucleic acids and proteins, is reportedly altered under diseased and infected states ^20-22^.

Macaques are relevant models of human pregnancy ^23^ and ZIKV infection during pregnancy ^24^. In the current study, a rhesus macaque TSC model was utilized as it allows for the study of trophoblast-type responses. Macaque TSCs derived from first trimester vCTB maintain cellular proliferation and can be directed to ST- or EVT-specific differentiation ^12^. We previously showed that TSCs differentiated to ST display features characteristic of early first-trimester, a developmentally critical period before the definitive placenta has completely formed. The objectives of this study were to 1) determine which macaque trophoblast cell types were permissive to ZIKV infection, 2) determine the molecular and secretory impact of ZIKV infection, and 3) to determine the utility of placental EVs (PEVs) to serve as readout of trophoblast infection status. The cellular and EV responses associated with infection presented here provide insight into how ZIKV impacts trophoblasts in the first trimester and suggests that PEV cargo may serve as a readout of placental ZIKV infection.

## Results

### Differentiation of TSCs alters cell permissiveness to ZIKV

Since maternal ZIKV infection early in pregnancy is associated with a high rate of pregnancy loss and fetal malformations, an early gestation in vitro macaque TSC model was used to determine which trophoblast cell types are permissive to infection. In vivo mouse ^25,26^ and porcine ^27^ studies and in vitro human studies ^28-30^ suggest that strains of the African lineage may be more detrimental to pregnancy than Asian lineage strains, hence, an African lineage strain was utilized to inoculate macaque TSCs. Preliminary time-course experiments to test multiplicities of infection (MOIs) were performed for each cell type to assess viral replication through 72 hrs of culture (Figure 1A). The amount of infectious virus detected in conditioned media from TSCs and syncytiotrophoblasts grown in suspension (ST3Ds) plateaued at 60 and 48 hrs, respectively. Inoculation with a MOI of 5 resulted in higher levels of infectious virus produced in TSCs compared to an MOI of 1 or 10, whereas viral replication was highest in ST3D’s with an MOI of 10.

**Figure 1.**
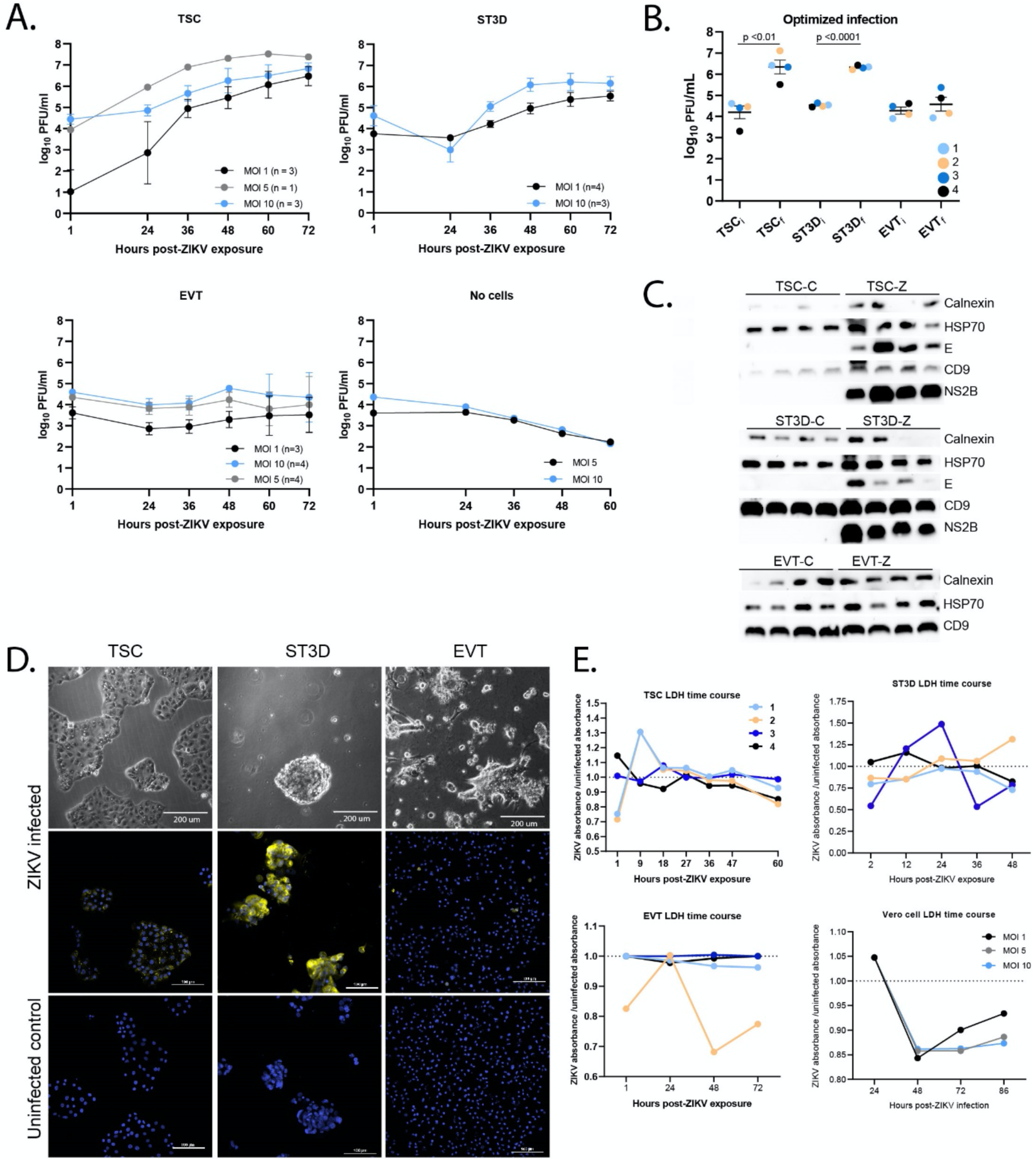
Early gestation trophoblast cells are permissive to ZIKV infection. **A)** Quantification of infectious virus by plaque assay in a time course and multiplicity of infection (MOI) dose response for each cell type (TSC, ST3D, EVT, or “no cells”). The number of cell lines tested, n, is indicated within parentheses next to the MOI. The quantity of infectious virus detected at each time point represents accumulated virus over time since the previous time point from the same well. **B)** Quantification of infectious virus by plaque assay with optimized MOI and duration of culture (i subscript = initial time point; f subscript = final time point) for each cell type and cell line (indicated by circle color). The mean plaque forming units (PFUs) ± the standard error of the mean (SEM) is shown. **C)** Western Blots of ZIKV E, NS2B, CD9, Calnexin, and Heat shock protein 70 (HSP70; loading control) proteins for each cell line (n = 4) and cell type. **D)** Immunostaining for ZIKV E protein. The top panel shows bright field images of trophoblasts after ZIKV exposure. ZIKV infected cells (middle panels) and control cells (bottom panels) were stained with an antibody against ZIKV E protein (yellow) and a nuclear stain (DAPI; blue). **E)** LDH time course data collected on TSCs, ST3Ds, EVTs, and Vero cells. Data is presented as the mean of biological (n=3) and technical replicates at each time point. The y-axis is presented as ZIKV LDH absorbance/control LDH absorbance, where a value > 1 indicates less cell death in ZIKV exposed cells and a value <1 indicates more cell death in ZIKV exposed cells.

An increase in infectious virus was not observed during the EVT time course. Since the quantity of virus did not decrease, to further verify viral replication, wells that did not contain cells but contained EVT culture extracellular matrix components (Matrigel and Col IV) (“no cells”) were exposed to virus and the media was evaluated by plaque assay. Compared to the amount of virus detected in the “no cells” wells at 60 hrs (mean 2.2 log_10_ plaque forming units (PFU)/ml), there was ∼40-fold more virus (mean 3.8 log_10_ PFU/ml) in the EVT samples at 60 hrs (Figure 1A). The infectious half-life of ZIKV is ∼12 hours ^31^, which indicates that the EVTs did indeed release a low level of infectious virus. For subsequent EVT infection experiments, cells were inoculated at an MOI of 5 and cultured for 72 hrs.

To minimize cytopathic effects and maximize information provided by EVs, the length of culture was chosen based on when the amount of virus being produced began to plateau, expecting that EVs released during peak viral shedding would show the most impact. MOIs for the optimized infection experiments were chosen based on whether there was a substantial increase in viral production (see methods). Of note, an initial (i) aliquot of culture medium was collected after ZIKV inoculation or mock inoculation and then a final (f) aliquot of medium was collected at the culture endpoint to assess viral replication by plaque assay (Supplemental Figure 1A). The quantity of infectious virus detected in TSC and ST3D media increased significantly by ∼100-fold between the initial and final time points within the respective cell type (TSC_i_ = mean 4.20 log_10_ ± SEM 0.16, TSC_f_= mean 6.35 log_10_ ± 0.17, ST3D_i_ = mean 4.53 log_10_ ± 0.04, ST3D_f_= mean 6.32 log_10_ ± 0.07; Figure 1B). A minimal change in infectious virus was detected in the EVT samples (EVTi = mean 4.28 log_10_ ± 0.14, EVT_f_= mean 4.58 log_10_ ± 0.23; Figure 1B) as anticipated from the data in Figure 1A.

Cellular ZIKV infection was validated by evaluating the presence of ZIKV E and NS2B proteins within inoculated and mock-inoculated trophoblasts. Both proteins were detected via Western Blot in the ZIKV exposed TSCs and ST3Ds but not in uninfected (control) cells (Figure 1C). Neither ZIKV protein was detectable in inoculated EVT samples (Supplemental Figure 3), supporting the minimal increase of infectious virus from the plaque assays. Positive ZIKV E protein detection via immunofluorescent staining further supported that TSC and ST3D are clearly permissive to ZIKV, while few EVTs stained positive for ZIKV E. (Figure 1D). Full Western Blot gel and isotype control immunofluorescence images are shown in Supplemental Figures 2 and 3, respectively.

To determine if the MOIs chosen for inoculation induced a cytopathic effect, cellular LDH was assayed in each cell type. Vero cells were used as a positive control as they are highly permissive to ZIKV, and cytopathic effects have been observed following ZIKV infection (Figure 1E). If ZIKV induced cell death, less LDH would be detected in infected cell media compared to uninfected control media resulting in a ZIKV:Control ratio less than one. A ratio of less than one was not consistently detected in trophoblast cultures, whereas a decrease in the LDH ratio was observed in Vero cells at 48 hrs that continued through 86 hrs. These data suggest that the MOI and viral strain chosen for this study did not result in trophoblast cell death during the culture period.

### ZIKV infection impacted cellular innate immune gene expression

RNA sequencing was performed to assess the impact of ZIKV on the cellular transcriptome. Transcriptomic analysis by poly(A)seq identified 82 (52 upregulated, 30 downregulated) and 113 (90 upregulated; 23 downregulated) differentially expressed genes (DEGs) in infected TSCs and ST3Ds, respectively, compared to controls (Figure 2A, 2B). Figure 2C presents a Venn diagram of upregulated mRNAs within the three trophoblast cell types. Transcriptomic analysis identified 13 genes significantly upregulated by ZIKV infection in both ST3Ds and TSCs (*APOL2, CXCL11, DDX58, IFIT2, IFNB1, IFNL1, INFL2, IFNL3, IRF1, ISG20, OASL, PARP14, and SCNN1D*). No significantly downregulated genes were similar between ST3Ds and TSCs. *PIANP* was the only gene found to be significantly upregulated (ZIKV mean 15.25 log_10_ ± SEM 2.72, control mean 0 log_10_ ± SEM 0) in EVTs exposed to ZIKV (EVT-Zs) (Figure 2D).

**Figure 2.**
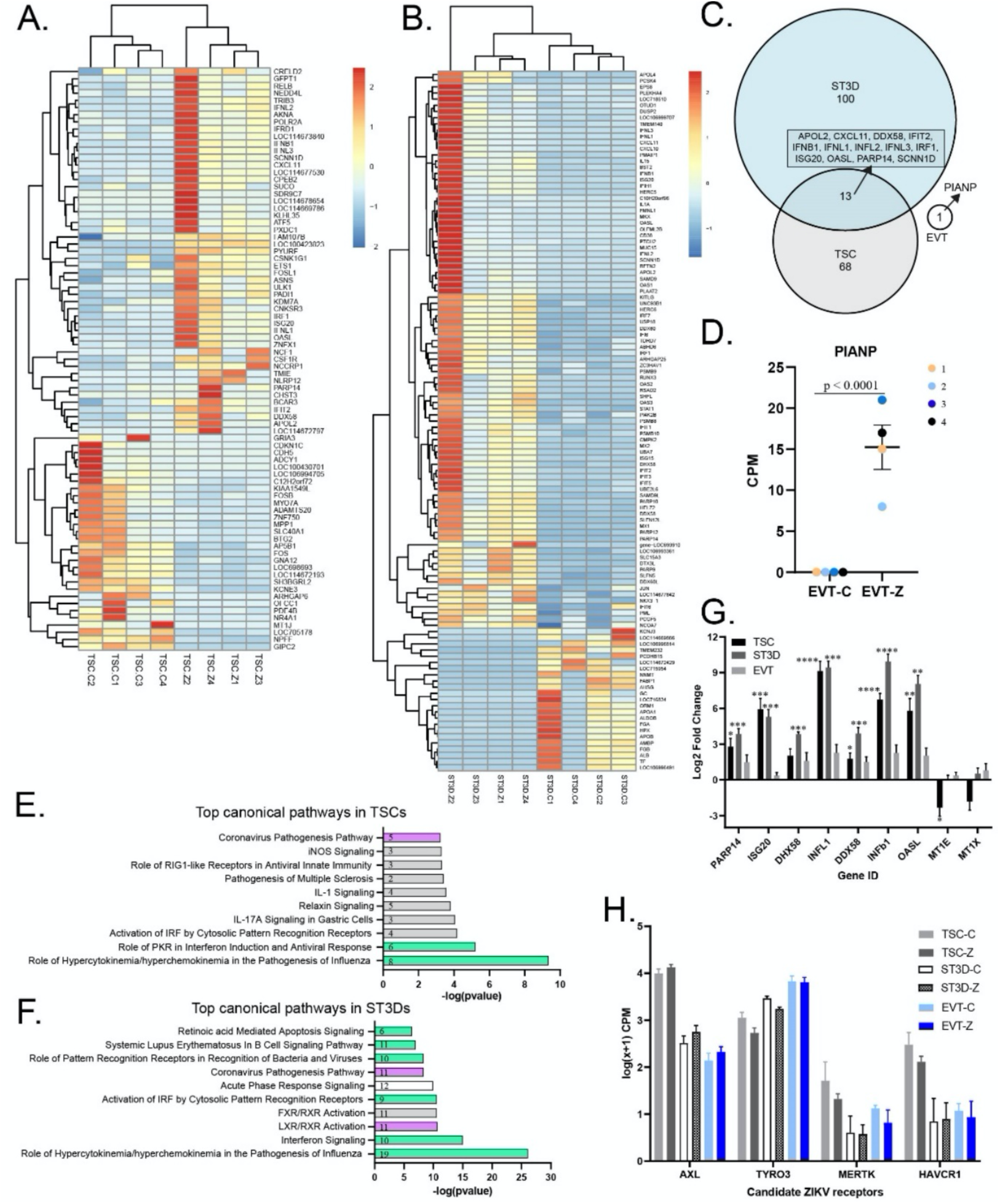
The impact of ZIKV on the cellular transcriptome. **A-B)** Heatmaps of significantly differentially expressed mRNAs in TSCs (**A**) and ST3Ds (**B**). **C)** A Venn diagram of the number of genes upregulated by ZIKV infection in TSCs, ST3Ds, and EVTs. **D)** The counts per million (CPM) of PIANP mRNA expression in EVTs with significance indicated on the graph. Replicate experiments are indicated by different colored dots. **E, F)**. Top 10 canonical pathways identified in the TSC and ST3D datasets (-log_10_ (p-value)). Purple indicates a positive Z-score (decreased in ZIKV); green indicates a negative Z-score (increased in ZIKV); white indicates 0 Z-score (no clear indication of whether the pathway is decreased or increased in ZIKV); and gray indicates unknown direction. The number of genes identified in each pathway is stated on the graph. **G)** A bar graph of differential mRNA expression (log_2_ fold change) validated by qRT-PCR. Significance is depicted on the graphs (**\*** p <0.05; ** p <0.01; *** p <0.001; **** p <0.0001). **H)** Normalized log (x + 1) transformed CPM determined by edgeR for candidate ZIKV receptors.

Integrated Pathway Analysis (IPA) was performed on the TSC (Figure 2E) and ST3D (Figure 2F) data sets. IPA identified “role of hypercytokinemia/hyperchemokinemia in the pathogenesis of Influenza” to be highly upregulated and the “coronavirus pathogenesis pathway” to be downregulated by ZIKV infection in TSCs and ST3Ds. The “role of PKR in interferon induction and antiviral response” was another top canonical pathway upregulated by ZIKV infection in TSCs (TSC-Z) (Figure 2E). Of the top five most significant diseases or functions identified by IPA in the TSC-Z cell mRNA data, four were related to viral replication and three were predicted to have decreased activation (data not shown).

The top canonical pathways upregulated by ZIKV infection in ST3D (ST3D-Z) cells included “interferon signaling”, “activation of IRF by cytosolic pattern recognition receptors”, “role of pattern recognition receptors in recognition of bacteria and viruses”, “systemic lupus erythematosus in B cell signaling pathway”, and “retinoic acid mediated apoptosis signaling” (Figure 2F). The “LXR/RXR activation” was downregulated in ST3D-Z cells. Diseases and function IPA of the ST3D-Z data predicts increased activation of the “antiviral response” and maturation of immune cells along with decreased replication of many viruses and “viral infection” (data not shown).

The expression of seven of the genes involved in the antiviral response pathway (*PARP14, ISG20, DHX58, INFL1, DDX58, IFNB1*, and *OASL*) and two genes involved in the anti-apoptosis pathway (*MT1E* and *MT1X*) ^32,33^ were validated with qRT-PCR (Figure 2G). The trends in expression observed with qRT-PCR agreed with those identified by poly(A)-seq analysis (Supplemental Table 3). TSC-Z or ST3D-Z samples had significantly increased expression of six and seven genes, respectively, with only minor elevations in *PAPR14, DHX58, IFNL1, DDX58, IFNB1*, and *OASL* genes in EVT-Z (Figure 2G). A trend in decreased expression of *MT1E* and *MT1X* was confirmed in TSC-Z samples by qRT-PCR, supporting the RNAseq data.

Potential receptors for ZIKV entry were detected in the Poly(A)seq data, including *AXL, TYRO3, MERTK*, and *HAVCR1* (*TIM1*) (Figure 2H) ^29^. *DC-SIGN* was not detected. *AXL* and *HACVR1* expression were greatest in TSCs whereas TYRO3 expression was greatest in EVTs. No significant difference in receptor expression was observed between infected and control cells.

### ZIKV infection altered the cellular miRNAome

Placental miRNAs are temporally expressed throughout gestation and have critical roles in trophoblast differentiation and function ^34,35^, thus the miRNAome was profiled to assess alterations in expression relative to infection. miRNA-seq identified expression of 326 miRNAs in TSCs, 335 miRNAs in ST3Ds, and 317 miRNAs in EVTs. In the TSCs, one miRNA (mml-miR-663) was significantly decreased and none were significantly increased in infected versus control cells (Figure 3A). Although ST3Ds were highly infected, no miRNAs were significantly impacted by ZIKV exposure. Despite modest productive infection of EVTs, 52 miRNAs were significantly increased and six significantly decreased in infected versus control EVTs (Figure 3B).

**Figure 3.**
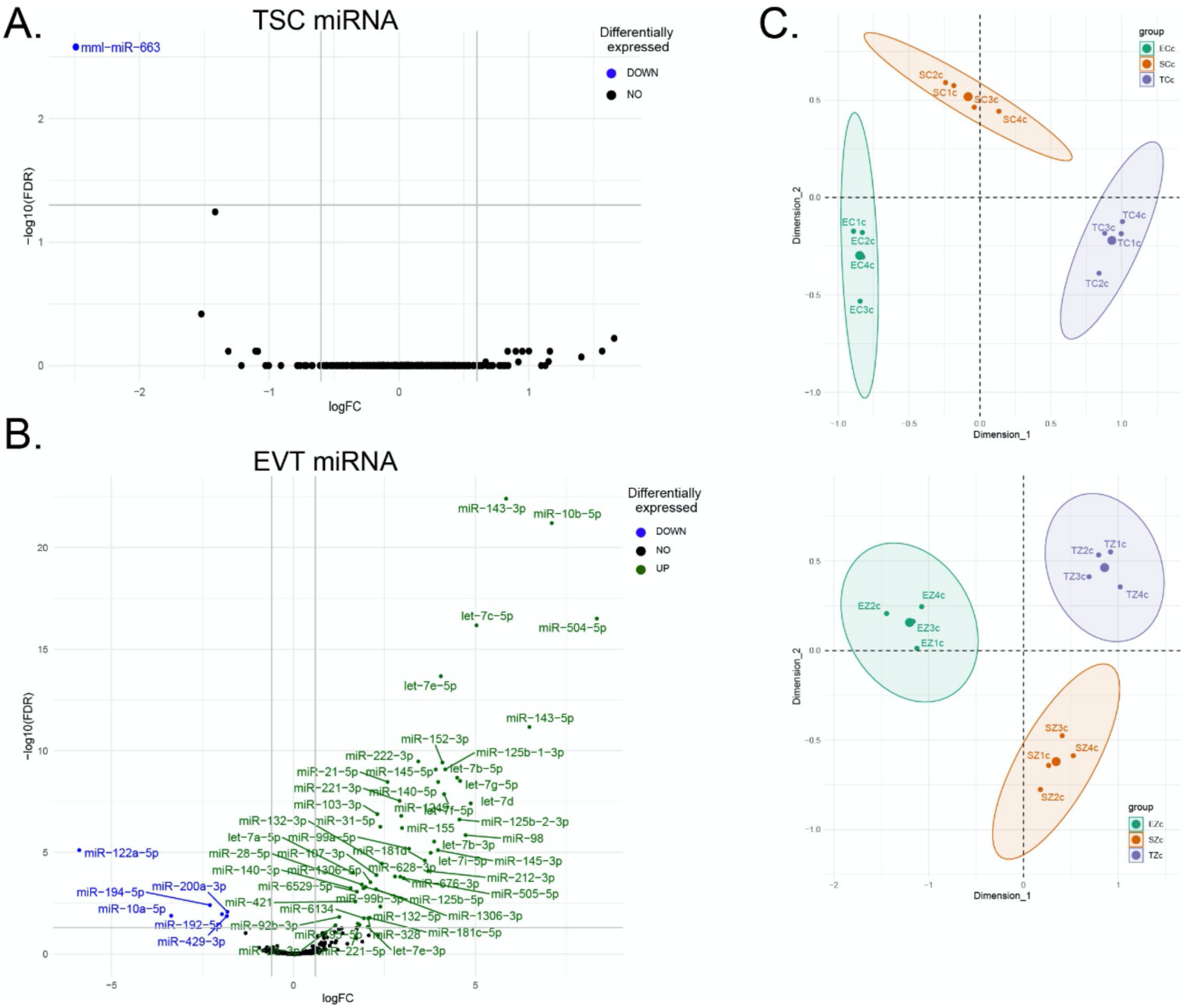
The impact of ZIKV exposure on the cellular miRNAome. **A-B)** Significantly differentially expressed miRNAs are displayed in color in the volcano plots for TSCs and EVTs (Blue indicates a decrease in ZIKV samples; Green indicates an increase; Black indicates no significant change). **C)** Multidimensional scaling (MDS) plots of miRNA expression across cell types for either control or ZIKV-exposed cells. The large dot indicates the center of the group.

To determine if ZIKV exposure resulted in similar trends of altered miRNA expression across trophoblast cell types, miRNAs significantly impacted in one cell type were compared to the other cell types (data not shown). Only miR-122a-5p trended towards decreased expression in all three cell types exposed to ZIKV (TSC 1.2 log_2_FC; ST3D 0.4 log_2_FC, and EVT 5.9 log_2_FC) with only a change in EVT expression statistically significant. Overall, cell types maintained a distinct miRNA profile regardless of ZIKV exposure (Figure 3C).

### ZIKV infection did not alter trophoblast secretion of hormones and modestly impacted cytokine secretion

To assess the impact of infection on trophoblast function, secretion of hormones, cytokines, chemokines, and growth factors within conditioned cell culture media was assayed. CG and progesterone are two key hormones in the recognition and maintenance of pregnancy. There were no differences in CG or progesterone secretion between infected and control samples for TSC, ST3D, or EVTs (Supplemental Figure 4A). TSCs secreted minimal CG or progesterone and ST3Ds and EVTs secreted significantly more CG than TSCs, in agreement with our previous report ^12^.

Cytokines and growth factors associated with infection and the inflammatory response were also quantified (Supplemental Figure 4B, 4C). *ITAC* (*CXCL11*) was significantly upregulated in infected ST3Ds compared to control (a similar trend was observed in *CXCL11* mRNA expression, Figure 2B), and was more highly expressed in infected ST3Ds compared to infected TSCs. Otherwise, no other significant differences in cytokine or growth factor secretion were observed between infected and control cells.

Regardless of infection status, there were significant differences in the secretion of several cytokines and growth factors between trophoblast cell types (Supplemental Figure 4B). ST3D and EVTs expressed significantly more *IL-1RA* and *bNGF* than TSCs. IL-6 was significantly upregulated in control EVTs compared to control ST3Ds. *FGF-2, VEGF-A, and VEGF-D* expression was significantly increased in ST3D cells compared to TSCs or EVTs. Conversely, some cytokines and growth factors showed no significant differences across cell type or infection status (Supplemental Figure 4C).

### Characterization and impact of ZIKV infection on EVs

Trophoblast-secreted EVs were isolated from ZIKV and control cell conditioned media to assess changes in their physical properties and cargo following ZIKV exposure. EV samples were characterized by Zetaview NTA (Figure 4A), and a consistent trend towards increased particle size was observed in TSC-Z and ST3D-Z EVs compared to their controls (Figure 4A). This trend also was seen with EVT samples, which were not widely infected. No differences in particle concentration were observed in any cell type, although Figure 4A shows that TSCs tended to release fewer EVs. TEM imaging verified the appropriate shape and size of isolated EVs (Figure 4B).

**Figure 4.**
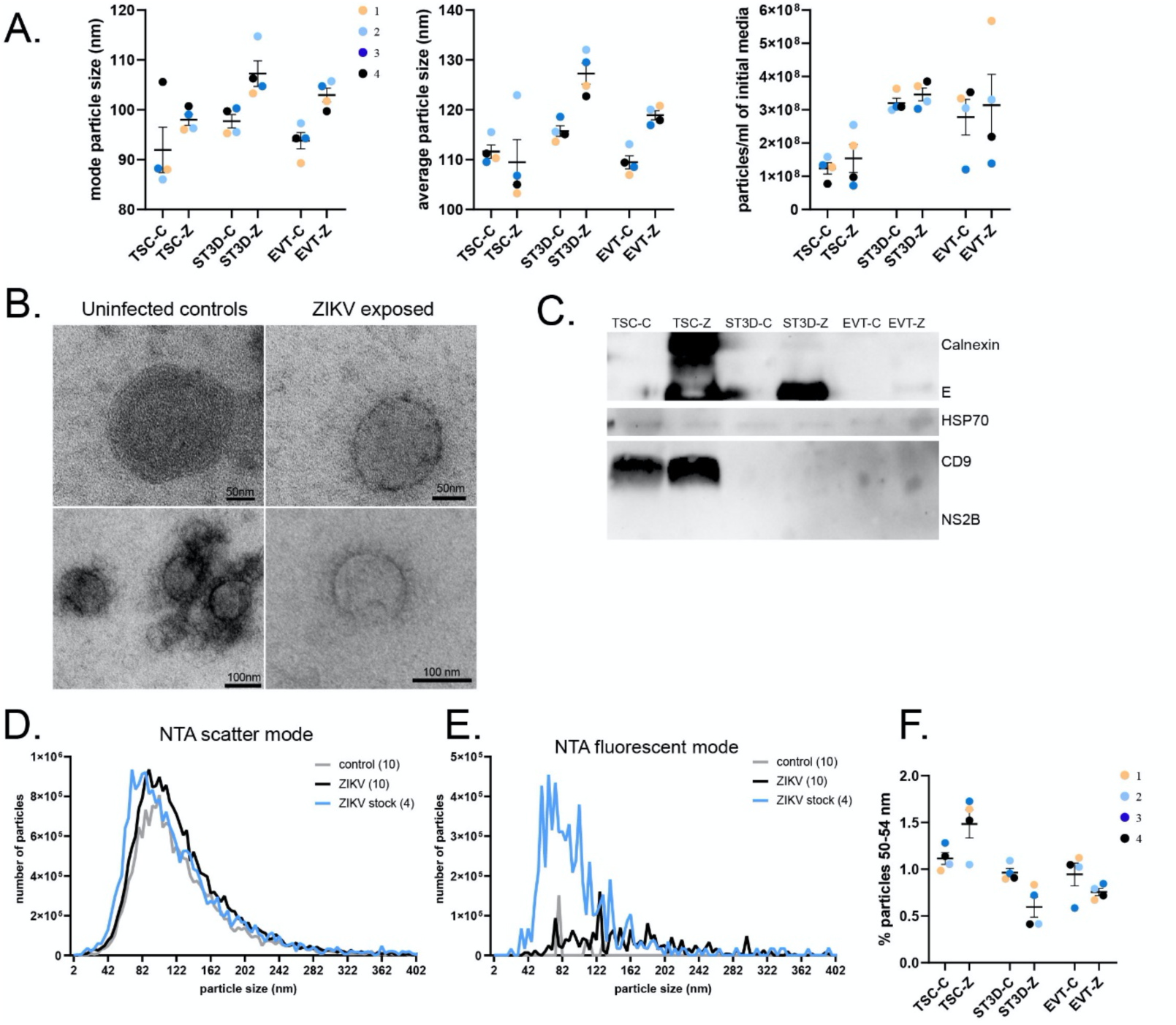
EV characterization from trophoblast-conditioned media. **A)** NTA analysis of the mode and average particle size in nm, and the concentration of particles presented as the particles/ml, on all 24 EV samples. **B)** Representative TEM images of four EV samples (two control and two ZIKV) with scale bars included. **C)** Western Blots of cell and EV samples for ZIKV E, ZIKV NS2B, Calnexin, HSP70, and CD9. **D-E)** EV samples were characterized with NTA scatter mode and fluorescent mode. The average number of particles detected across the control samples, ZIKV samples, and ZIKV stock were plotted. The number of samples is indicated in the parentheses. **F)** The percentage of “ZIKV-sized” particles (50-54 nm) was calculated for all 24 EV samples and is plotted by the treatment group.

To determine if ZIKV proteins were packaged into EVs, EV preparations were assessed for E and NS2B proteins by Western Blot. The ZIKV E protein was readily detected in the TSC-Z and ST3D-Z EV samples with little detected in the EVT-Z EV samples and none in any controls (Figure 4C; Supplemental Figure 3D). Since EVTs were modestly infected with ZIKV, it is not surprising that the E protein was not detected. The presence of two proteins commonly enriched in EVs, CD9 and HSP70, was also determined. On this blot CD9 was only detected in the TSC-C and TSC-Z EVs; however, on another blot, EV samples (those submitted for mass spectrometry) from all three cell types were positive for CD9 (Supplemental Figure 3E). Calnexin, an ER resident protein not expected to be present within EVs, was confirmed to be absent from all EV samples, and, as expected, calnexin was observed in cell samples along with CD9 (Figure 1C). Interestingly, a higher molecular weight cross-reactive band was seen in the TSC-Z sample. Calnexin was not detected in EVs by mass spectroscopy so it is unlikely that the cross-reactive protein is calnexin; also of note is that this band is slightly larger than that observed in the cell lysates (Supplemental Figure 3D).

To assess whether the ZIKV E protein associated with EVs, EVs were stained with a fluorescently labeled ZIKV E antibody and analyzed by Zetaview NTA. EV preparations and concentrated ZIKV stock (positive control) were analyzed under scatter and fluorescent modes (Figure 4D and 4E, respectively). The average number of particles at a given particle size were calculated and are depicted as a single histogram. Zika virions are ∼ 50 nm in size ^36^ and a shift to the left was observed in the “ZIKV stock” histogram compared to the EV preparations, indicative of the abundant presence of smaller particles which are potentially ZIKV virions (Figure 4D) (larger particles in this preparation may represent Vero cell-secreted EVs). The fluorescence histogram of EVs isolated from ZIKV cultured trophoblasts, termed “ZIKV” in the figure, showed that the size of particles ranged from ∼50-300 nm (Figure 4E). The detection of ZIKV E protein in particles larger in size than virions suggests that the E protein was associated with EVs. Of note, minimal background fluorescence was detected in the control EV samples. To further support these data, the number of particles detected in the size range of ZIKV (∼50 nm) were quantified (reanalysis of the 24 samples shown in Figure 4A) and the percentage of particles between 50 and 54 nm was calculated (Figure 4F). ZIKV ST3D and EVT EV samples contained fewer “ZIKV-sized” particles than controls, again suggesting that Zika virions were not abundant in the EV samples. Overall, these data indicate that ZIKV E protein is a component of EVs.

DAVID enrichment analysis was used to further verify the mass spectroscopy and Poly(A)-seq data gathered from EVs (Figure 5A and 5B). Proteins identified among all 24 EV samples (ZIKV and control) were pooled and analyzed with DAVID. The mass spectrometry results showed remarkably high enrichment of the “extracellular exosome” cellular component (Figure 5A). Other highly enriched cellular components included focal adhesion, as well as membrane, plasma membrane, and cytosol, as expected. Based on the top 1000 genes identified in the eight EV poly(A)-seq samples, the most enriched cellular component also was “extracellular exosome” (Figure 5B). Lastly, IPA on all proteins detected in the TSC, ST3D, and EVT EV mass spectroscopy data show that ∼50% of proteins isolated were cytoplasmic with ∼20-25% associated with the plasma membrane across all cell types (Figure 5C). Enzymes, transporter proteins, kinases, transcription regulators, translation regulators, transmembrane receptors, and peptidases were some of the more abundantly detected protein types (Figure 5D). Altogether, these data support the authenticity of these trophoblast-produced EV preparations.

**Figure 5.**
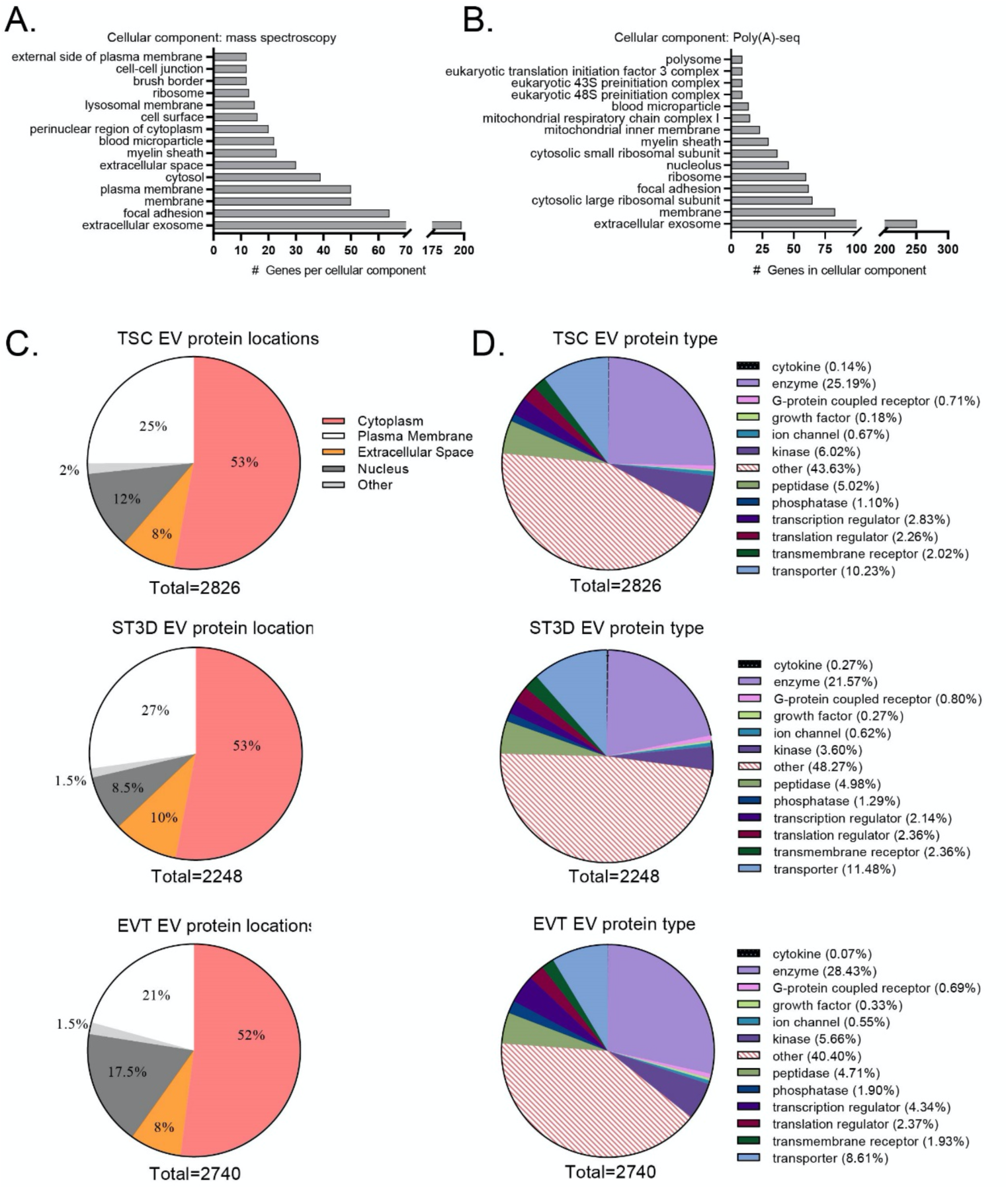
EV verification and protein information analyses. **A)** DAVID pathways enrichment on the 392 proteins identified in all 24 EV samples. **B)** DAVID pathways enriched on the top 1,000 genes detected among the eight EV samples by poly(A)-seq. For DAVID enrichment analysis, a minimum of five genes for that pathway/process and an adjusted p-value < 0.05 (Benjamini-Hochberg correction) were required. **C-D)** IPA on all proteins detected in the TSC, ST3D, and EVT EV data provide the protein location and type.

### Mass spectroscopy analysis on EV proteins

Proteomic data analysis revealed that many proteins were significantly differentially detected in TSC (191 total: 180 up, 11 down), ST3D (75 total: 63 up, 12 down), and EVT (128 total: 78 up, 50 down) EVs released from ZIKV-exposed cells (Figure 6A-C). Interestingly, unsupervised clustering of protein expression revealed that samples do not group by infection status.

**Figure 6.**
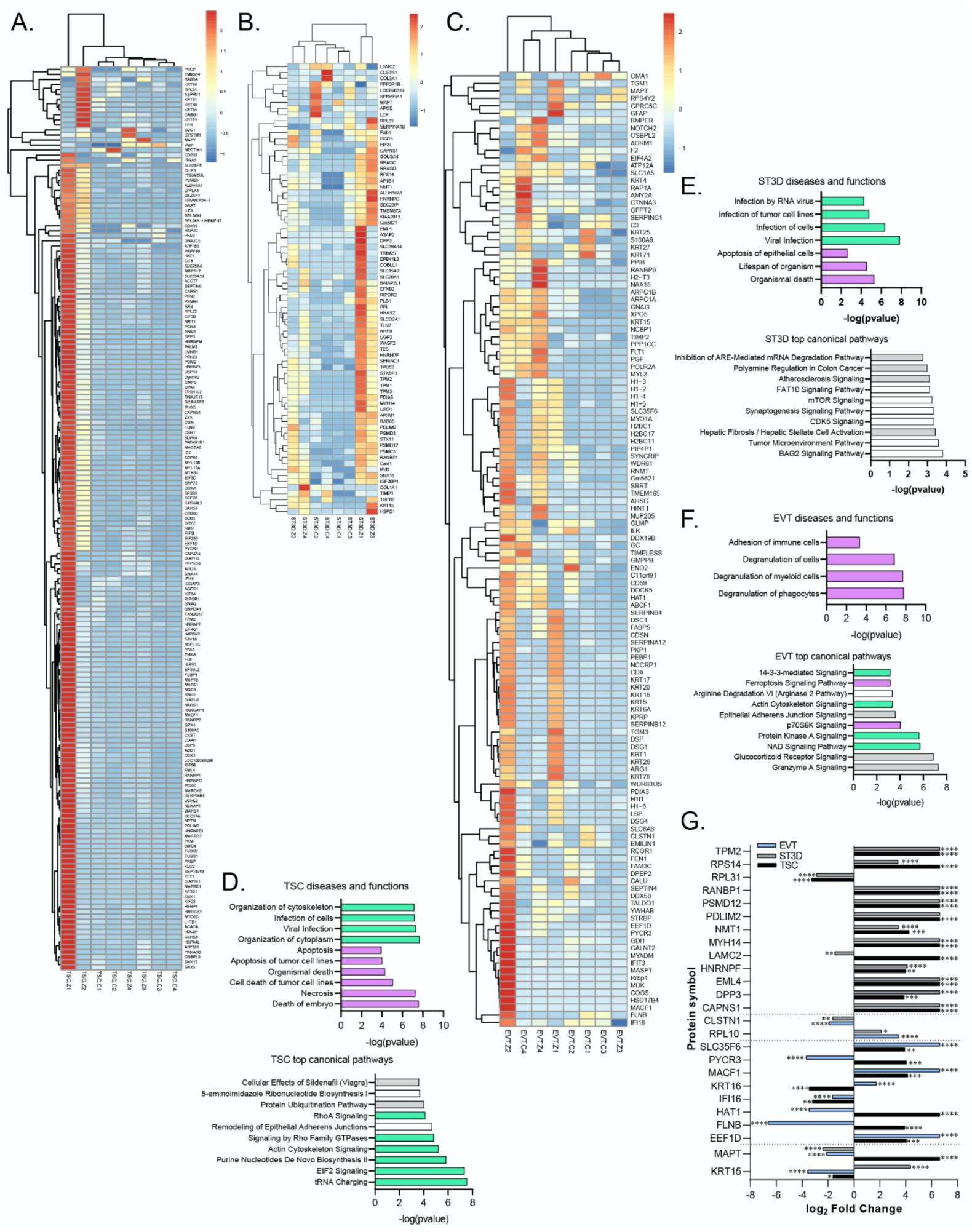
EV proteomics analysis. **A-C)** Heatmaps of significantly differentially detected proteins in TSC, ST3D, and EVT EV samples. **D-F upper graphs)** Top diseases and functions identified in the TSC, ST3D, and EVT EVs. **D-F lower graphs)** Top canonical pathways identified in the TSC, ST3D, and EVT EVs. Purple indicates a positive Z-score (predicted decreased in ZIKV); green indicates a negative Z-score (predicted increased in ZIKV); white indicates 0 Z-score (no clear indication of whether the pathway is decreased or increased in ZIKV); and gray indicates unknown direction. **G)** Log_2_ fold change of genes identified in multiple cell types. A positive log_2_ fold change indicates increased detection in ZIKV EVs and negative indicates decreased detection in ZIKV EVs. Significance is depicted on the graphs (**\*** p <0.05; ** p <0.01; *** p <0.001; **** p <0.0001).

IPA was performed to predict the disease and biological functions and canonical pathways that the EV proteins may have a role in. Top diseases and biological functions predicted for TSC-Z EV proteins included decreased functions associated with death (necrosis, apoptosis) as well as increased cellular organization and infection (Figure 6D upper graph). The topmost significant predicted canonical pathways increased in TSC-Z EVs were tRNA charging, EIF2 signaling, purine nucleotide biosynthesis, actin cytoskeleton signaling, Rho GTPase signaling, and RhoA signaling (Figure 6D lower graph). In comparison, ST3D-Z EV proteins were associated with decreased death, lifespan, and apoptosis with enrichment for infection and viral infection (Figure 6E upper graph), and ZIKV infection was predicted to impact various other pathways but directional shifts were not identified (Figure 6E lower graph). Decreased degranulation of various cells and decreased adhesion of immune cells were predicted for EVT-Z EV proteins (Figure 6F upper graph). EVT-Z EVs had increased NAD, Protein Kinase A, actin cytoskeleton, and 14-3-3 mediated signaling, whereas ferroptosis and p70S6K signaling pathways were decreased (Figure 6F lower graph).

To determine if EV protein cargo was shared among the three cell types, proteins significantly differentially detected by mass spectroscopy were compared. This comparison revealed two proteins common among the three cell types, eight proteins between TSC and EVT, two proteins between ST3D and EVT, and 13 proteins between TSC and ST3D (Figure 6G). The log_2_ fold change is plotted for each of those proteins with a positive change indicative of increased presence in ZIKV EVs and negative change indicative of decreased presence in ZIKV EVs. TPM2, RPS14, RANBP1, PSMD12, PDLIM2, NMT1, MYH14, HNRNPF, EML4, DPP3, CAPNS1, RPL10, SLC35F6, MACF1, and EEF1D were increased in ZIKV EVs. RPL31, CLSTN1, and IFI16 were consistently decreased in ZIKV-EVs. The other six genes (LAMC2, PYCR3, KRT16, HAT1, FLNB, MAPT, KRT15) were increased in some cell types but decreased in others.

In terms of placenta specific protein detection in EVs, a macaque-specific placental MHC class I molecule, MAMU-AG4 (Uniprot ID: Q5TM80) was detected in all 24 samples. Endogenous retrovirus group FRD member 1 (Uniprot ID: A0A1D5R0I7) was detected in six of eight ST3D EV samples. Placenta growth factor (Uniprot ID: A0A1D5R9B1) and pregnancy specific beta-1-glycoprotein 7 (Uniprot ID: F6YVT1) were detected in six of eight EVT EV samples. Pappalysin 1 (Uniprot ID: F6ZUN7) was detected in all ST3D and EVT EVs but no TSC EVs.

### The impact of ZIKV on EV poly(A) and miRNA cargo

A total of 24 EV samples were submitted for Poly(A)-seq but cDNA libraries could only be prepared for eight samples (3 TSC-C, 1 TSC-Z, 2 ST3D-C, and 2 ST3D-Z). A total of 31,459 transcripts were identified of which 2,618 transcripts were detected in all eight EV samples. Conversely, 3,898 and 9,490 transcripts were specific to either ZIKV or control EV samples, respectively. When comparison was restricted to transcripts detected in a majority of samples regardless of cell type origin (3/5 control or 2/3 ZIKV-exposed; Figure 7A), 832 and 601 transcripts were detected only in control or ZIKV EVs, respectively. A total of 69 and 41 transcripts were identified in all three ZIKV EV samples or all five control EV samples, respectively. Immune response related genes were detected in EVs released by ZIKV-infected cells. IFNL1 transcript XM_015123826.2 was only detected in EVs released by ZIKV-infected cells. Two different IL1RN isoforms were detected in the control or ZIKV EVs, with each transcript being unique to either group (XM_001091833.4 and XM_015113137.2 in control or ZIKV, respectively. Pathways enriched by transcripts identified in the majority of ZIKV or control EVs (602 and 871 for ZIKV and control, respectively) are shown in Figure 7B.

**Figure 7.**
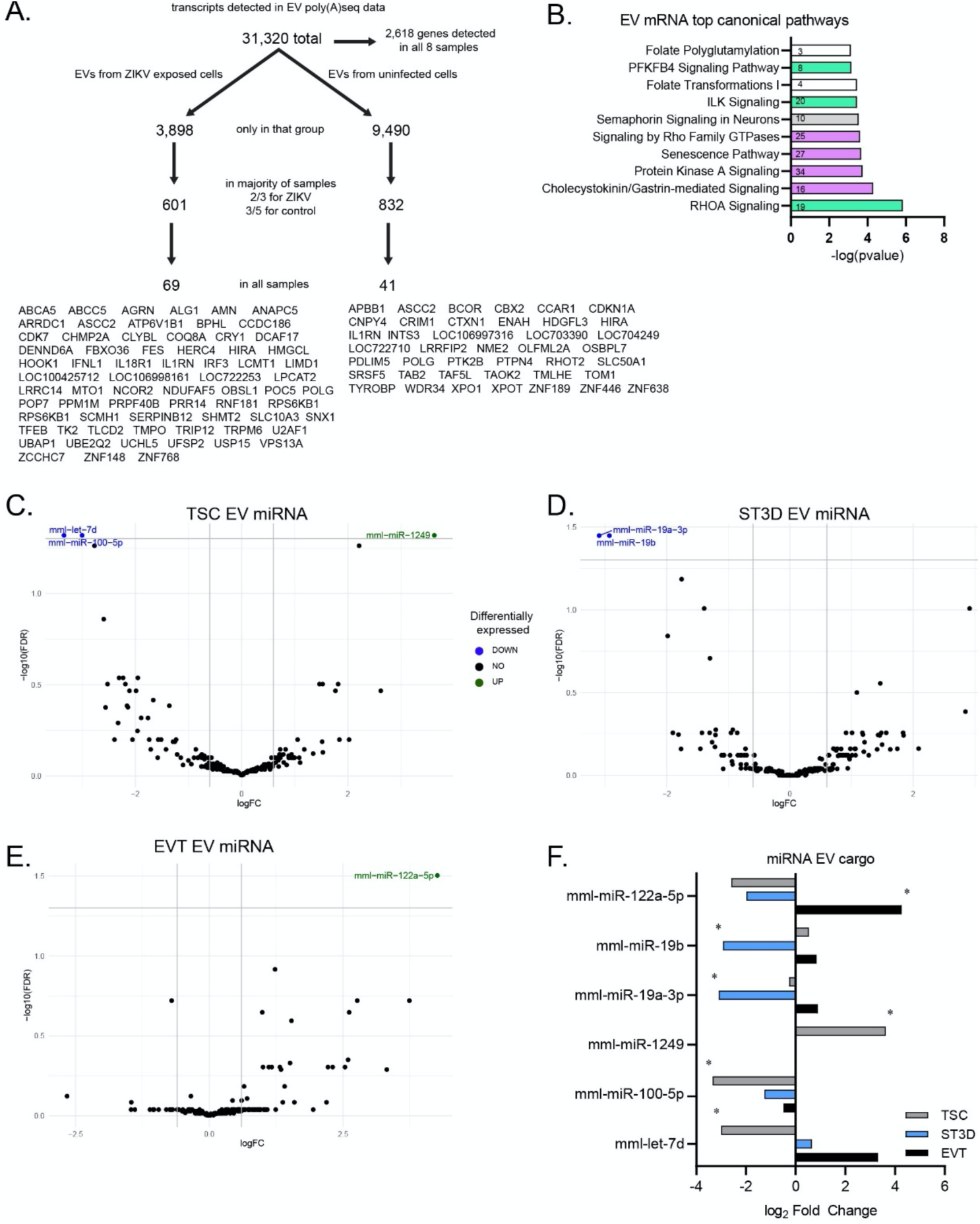
ZIKV exposure impact on EV mRNA and miRNA cargo. **A)** Schematic of the total number of transcripts detected across the eight EV samples, the number of transcripts solely detected in either the ZIKV or control samples, the number of transcripts detected in a majority (three of five control; two of three ZIKV samples) of the samples per group, and the number of transcripts detected in all samples of that group. The 66 and 35 gene IDs that correspond with transcripts identified are presented below. **B)** Top 10 canonical pathways identified in a majority of either the ZIKV or control EV samples (-log_10_ (p-value)). Purple indicates a positive Z-score (decreased in ZIKV); green indicates a negative Z-score (increased in ZIKV); and white indicates 0 z-score (no clear indication of whether the pathway is decreased or increased in ZIKV). The number of transcripts identified in each pathway are stated on the graph. **C-E)** Volcano plots of miRNAs detected in TSC, ST3D, and EVT EVs. miRNAs significantly differentially expressed are highlighted on the plots (blue = down in ZIKV exposed; green = up in ZIKV exposed; black = not significantly different). **F)** A bar plot depicting the log2 fold change for the six miRNAs found to be significantly different in EVs released by at least one of the cell types. * indicates an FDR <0.05.

### miRNA-seq on EVs

A total of three miRNAs were detected at significantly different levels between ZIKV exposed and control TSC EVs (Figure 7C). mml-miR-1249 was significantly increased by ZIKV exposure and mml-miR-100-5p and mml-let-7d were significantly decreased. mml-miR-19b and mml-miR-19a-3p were significantly decreased in the ST3D-Z EV samples while none were increased (Figure 7D). Only mml-miR-122a-5p was significantly increased in the EVT-Z EVs and none were significantly decreased (Figure 7E).

To determine if ZIKV exposure resulted in similar trends of altered EV miRNA cargo across different cell types, miRNAs significantly impacted in one cell type were compared to the other cell types (Figure 7F). mml-miR-100-5p was decreased in all three cell types exposed to ZIKV and was the only miRNA with similar trends observed among the three different cell types. miR-1249 was only detected in EVs released by TSCs. mml-let-7d was significantly decreased in EVs released by TSC-Z and almost significantly increased in EVs released by EVT-Z (FDR = 0.51).

## Discussion

To determine which trophoblast cells are permissive to ZIKV infection and to understand how infection impacts function, we established a novel in vitro primate trophoblast cell model. TSCs and ST3Ds were highly permissive to infection as indicated by the presence of infectious virus in the culture medium and ZIKV proteins detected within cells. In addition, disease and function IPA on the TSC an ST3D mRNA data indicate these cells responded to the ZIKV exposure. Conversely, EVTs maintained a level of resistance to ZIKV infection as evident by low virus production. ZIKV exposure also impacted EV size and cargo, including detection of the ZIKV E protein. Thus, experimental infection of in vitro macaque trophoblasts presented in this study supports the hypothesis that ZIKV can infect early gestation trophoblast cells. These results are consistent with reports of trophoblast permissiveness to ZIKV infection in human first trimester trophoblast cells, first trimester placental explants, and embryonic stem cell (ESC)-derived trophoblast cells ^29,37,38^. The model used here supports that early gestation trophoblasts are permissive to ZIKV infection and that the impact of placental cell infection is reflected in the composition of trophoblast EVs.

ZIKV exposure induced expression of genes involved in a cellular immune response. Cells maintain multiple mechanisms to identify and curtail a viral infection, including inhibiting protein translation, degrading viral mRNA transcripts, and alerting immune cells to infection via proinflammatory cytokine secretion ^39,40^. The protective effect of IFNs against ZIKV infection has been previously reported ^41-43^. Elevated gene expression of IFNs and IFN pathway-associated genes in the current study was noted, which agrees with previous studies ^43-45^. Human ESCs-derived trophoblasts, which are described as primitive trophoblasts, expressed low levels of type I and III IFNs ^29^. However, in the current study IFN-beta (type I IFN) and IFNL1 and IFL3 (type III IFNs) transcript abundance increased after ZIKV exposure. Provocatively, IFNL1 mRNA was detected in EVs from ZIKV exposed cells but not control. The different responses between human ESC-derived trophoblasts and the rhesus TSC-derived trophoblasts in the current study suggests these two cell types represent different developmental stages. It should be noted that no changes in IFN concentrations measured by Luminex assay were detected in response to ZIKV infection and is potentially due to either post-transcriptional mRNA degradation or suboptimal cross-reactivity or sensitivity of the Luminex assay with the macaque proteins.

miRNAs also are important for trophoblast antiviral responses ^46^. Interestingly, the predominant changes in miRNA expression were observed in EVTs, the cell type that appeared to control/limit ZIKV infection compared to TSCs or ST3Ds. This raises questions as to whether these cells were able to respond more readily and whether changes in miRNA expression were involved in controlling the infection. mml-miR-122a-5p was decreased in all three trophoblast cell types. Interestingly, EVT-Zs had significantly decreased quantities of cellular mml-miR-122a-5p, while significantly increased quantities of the miRNA were detected in EVs of EVT-Z. mml-miR-122a-5p inhibits cellular proliferation, migration, and invasion in Pancreatic ductal adenocarcinoma cells^47^. Although the functional significance in trophoblast function remains to be investigated, the consistent trend towards decreased presence of mml-miR-122a-5p in the three trophoblast cell types raises questions as to its role in viral infection.

Despite changes in the miRNAome and transcriptome, the EVTs displayed modest levels of ZIKV replication, which is unlike findings with other in vitro models ^37,38,48^. The low level of EVT infection in the current study is an important finding as endovascular EVTs that migrate into and remodel maternal spiral arteries are in direct contact with maternal blood, circulating cells of the maternal immune system, and potentially ZIKV within maternal circulation. One possible explanation for the lack of ZIKV replication in our EVT cultures may be due to the presence of extracellular matrix components (collagen IV and Matrigel). Experimental infection of cancer cell lines with herpes simplex virus type 1 in the presence of Matrigel resulted in decreased viral replication, and the authors concluded that the extracellular matrix hindered viral replication post-viral entry ^49^. Another explanation could be the lack of ZIKV receptors. Although the presence of candidate ZIKV receptors was lower in EVTs compared to TSCs or ST3Ds, their presence, along with detection of two ligands potentially involved in ZIKV entry (GAS6 and PROS1) ^29^, suggests ZIKV could at least bind to these cells, an interpretation borne out by impact on EVT-Z EV cargo. The point at which EVTs can control infection remains unclear and future studies are needed to fully elucidate their response to ZIKV exposure.

Analysis of placental EV proteins, mRNAs and miRNAs revealed a range of cargo. For both mass spectrometry and poly(A)-seq, genes associated with extracellular exosomes were the predominant cellular component identified. Other proteins identified by mass spectrometry were associated with focal adhesion, plasma membranes and cytosol, and protein locations were predominantly the cytoplasm and plasma membrane. Poly(A)-seq also identified transcripts for many genes involved in translation, including ribosomal subunits and preinitiation complex factors. Another study found translational machinery within EVs ^50^ and the data herein support this previous observation.

Ultimately, the identification of biomarkers from circulating EVs could allow insight into placental health, or in the context of ZIKV, placental infection. Several placenta-specific proteins were identified in EVs from ST3Ds and EVTs, validating their placental cell origin. Specifically, Mamu-AG, an MHC class I molecule highly expressed in the rhesus placenta, was detected in all samples. Putative biomarkers identified in this study include the detection of ZIKV protein in EVs in addition to changes in EV cargo. The overlap between the ZIKV life cycle and exosome formation ^51,52^ could allow for Zika viral proteins, genomes, and/or whole virions to be packaged within exosomes. Here, EVs within the exosome size range stained positive for the ZIKV E protein by NTA, and ZIKV E protein but not the NS2B protein were detected in EVs by Western Blot. This is an important observation, as the NS2B protein is not packaged in the mature virion whereas the E protein is present. Although a low abundance of virions in the EV samples cannot be ruled out, the size of the ZIKV-E positive particles suggests incorporation of the E protein into EVs.

Two putative biomarkers of placental cell infection identified in EVs from ZIKV-exposed cells include Proteasome 26S subunit, non-ATPase 12 (PSMD12) and PDZ LIM domain protein 2 (PDLIM2). PSMD12 is involved in normal removal of damaged/misfolded proteins ^53^. In this context, PSMD12 packaged into EVs may chaperone damaged proteins into the secretory pathway for degradation in lysosomes. Since exosomes are a part of the secretory/endosome pathway, the increased presence of the PSMD12 in ZIKV-EVs could be part of a cellular immune response to increase degradation of viral proteins. Future studies should determine if functional PSMD12 is taken up by recipient cells. PDLIM2 is a ubiquitin E3 ligase that targets STAT2 for degradation ^54^ and in response to infection with other Flaviviridae viruses becomes upregulated^54^. In this study, PDLIM2 was increased in TSC-Z and ST3D-Z EVs with no change in cellular expression. One interpretation is that infected cells secreted additional PDLIM2 in an attempt to hinder the immune response of other cells to ZIKV. Additional studies are needed to better understand why and how PDLIM2 ends up in EVs and what impact this could have on recipient cells.

Overall, this study shows that macaque trophoblasts representative of early first trimester trophoblasts are permissive to ZIKV infection. STs are in direct contact with maternal blood and therefore could be readily infected. Although the location and presence of the TSC niche in the human or primate placenta is poorly understood ^55^, these cells are not in direct contact with maternal blood but could become infected if the ST layer is breached. As a progenitor cell that will differentiate to vCTB or column cytotrophoblasts to then give rise to ST and EVTs ^56^, their permissiveness to ZIKV could have major implications on placental health. In addition, we saw that infection altered trophoblast function, even in EVTs that did not show prominent infection. ZIKV exposure also altered EV composition and EVs released by infected cells may contain ZIKV proteins. Altogether, these changes could impact the maternal response to the pregnancy and maternal immune response to ZIKV at the maternal-fetal interface. Furthermore, alterations in EV composition could be used to noninvasively identify placental ZIKV infection, which could be of significant clinical importance, not only for ZIKV but other TORCHZ infections.

## Methods

### Trophoblast cell culture

Rhesus monkey TSCs ^12^ were graciously provided by Dr. Jenna Kropp Schmidt. The four TSC lines used for this study were: rh010319 (male; gestation date (gd) 58; line 1); rh090419 (female; gd 75; line 2), r121118 (male; gd 62; line 3), and rh020119 (female; gd 74; line 4). Previously published protocols ^12^ were followed with some minor modifications as described. TSCs were differentiated into either EVTs or ST3D aggregates (Supplemental Figure 1A). TSCs were grown in 13 ml media, EVTs in 15 ml media, and ST3Ds in 10 ml media (Supp Figure 1B).

TSCs were differentiated to EVT by seeding into T75 flasks coated with 1 μg/ml collagen-IV (col IV; Corning, NY, USA, Cat #354233) with 2% Matrigel (Corning, Cat # 354234) added to the EVT media. No additional Matrigel was added on day 3 when the media were changed as previously described. Growth Factor Reduced Matrigel (0.5%; Corning, Cat #354230) was added on day 6.

For ST3D culture, TSCs were plated in non-adherent T75 flasks (Thermo Fisher, Cat #174952 and Cat #156800) and cultured for 5 days total. Three days after initial plating, the media were removed, and fresh media added. For additional details on the media composition and growth conditions, please see Schmidt, et al. 2020 ^12^.

### ZIKV stock

ZIKV, DAK AR 41524, NR-50338 (“DAKAR”) was obtained through BEI Resources, NIAID, NIH, as part of the WRCEVA program. To propagate this stock for subsequent experiments, the previously published protocol was followed ^57^ with minor adaptations. Briefly, Vero cells (ATCC, Manassas, VA, USA, Cat #CCL-81) were plated at 4 × 10^6^ cells/T75 flask in MEM (ThermoFisher, Cat #11-095-080) containing 2% FBS (Peak Serum, Wellington, Colorado, USA, Cat #PS-FB1) and 100 mM sodium pyruvate (Sigma Aldrich, St. Louis, MO, USA, Cat #2256). The following day, they were exposed to DAKAR at an MOI of 0.1 for 1 hr in 2 ml of MEM medium. Cytopathic effect was observed at 48 hrs in the ZIKV exposed cultures, and media were collected and spun at 15,000 × g for 30 mins at 4°C. The supernatants were aliquoted and frozen at -80°C until used for infection studies. The final stock was sequenced by Dr. Shelby O’Connor’s laboratory at the University of Wisconsin-Madison using an Illumina MiSeq instrument and compared to the ZIKV stock reference sequence (Genbank Accession KY348860). Three single nucleotide (nt) substitutions at nt positions 470 (synonymous mutation; serine -> serine), 3868 (missense, alanine - valine), and 3790 (missense, alanine - valine) were identified in this ZIKV stock. The frequencies of these substitutions were all below 16%. The stock concentration (4.6 × 10^7^ PFU/ml) was determined by plaque assays that were run in triplicate, as previously described ^58^.

### Trophoblast ZIKV Inoculations

Duration of culture and MOI were first optimized prior to generating experimental infection replicates (Figure 1A). Both ZIKV-infected cells and uninfected controls underwent the same processing except uninfected cells were exposed to ZIKV-free Vero cell conditioned media that was collected alongside the ZIKV stock (mock infected). The MOI used was determined based on the quantity of virus calculated from the Vero cell plaque assay described above.

For infection of the TSCs and EVTs, one flask of cells from each line was lifted and counted just prior to infection. For TSCs and EVTs, media were aspirated, and 1 ml of inoculum was added to the flask. Cultures were incubated at 37°C and rocked gently every 15 mins for 1 hr. The inoculum was removed, and the cells were washed once with PBS followed by addition of fresh medium. The number of ST3D “cells” (at this point they are aggregates) present was based on the number of TSCs added to the flask three days prior (Supplemental Figure 1B). To infect the ST3Ds that were grown in suspension, aggregates/cells were pelleted by centrifugation at 500 × g for 3 mins, the supernatant was removed, and the cells were resuspended in 300 μl of inoculum. Aggregates were incubated in the ZIKV inoculum for 2 hrs at 37°C and gently every 30 mins. Since ST3Ds are grown in suspension, a larger volume of inoculum was used, and the length of inoculation was extended to account for this increase. Next, 2 ml of PBS was added, the cells were re-pelleted by centrifugation as above, and the supernatant was removed. A volume of 10 ml ST3D medium was added to the cells, and they were then transferred back into a T75 flask.

Once fresh media were added back for all cell types, a 500 μl aliquot was immediately removed to serve as a baseline of initial viral titer in the culture. The aliquot was spun at 500 × g for 5 mins to remove any cellular debris and a plaque assay was conducted. For EVTs, Growth Factor Reduced Matrigel (0.5%; Corning, Cat #354230) was supplemented to the cultures after this step. EVTs were cultured for a day longer than previously reported ^12^ (72 hrs total) to extend the duration of ZIKV infection. An MOI of 5, 10, and 5 were used to infect the TSCs, ST3Ds, and EVTs, respectively. Inoculated and mock-inoculated control TSCs were cultured for 60 hrs, while ST3Ds and EVTs were cultured for 48 and 72 hrs, respectively.

### Sample collection: cells and media

At the end of the culture period, TSCs were lifted, EVTs were scraped off the flasks, and ST3Ds were pelleted prior to aliquoting cells for RNA, DNA, and protein isolation (Supplemental Figure 1A). Conditioned media collected from all flasks were spun at 500 × g for 5 mins to remove dead cells and debris, pooled, and then aliquoted and frozen back at -80°C for EV isolation, hormone analysis, Luminex assay, or plaque assay.

### EV isolation

To isolate EVs, 20 ml of medium was placed onto a concentration column (Vivaspin 20; Sartorius, Swedesboro, NJ, USA, Cat # 1208L91) and spun for 60 mins at 3,000 × g at room temperature (RT). For EVT EV samples, due to a high density of Matrigel in the conditioned media, the media were first filtered using a 0.22 um filter (Millipore Sigma, Cat # SLGP033RS) and then spun for 60 - 90 mins. The concentrated sample was passed through a size exclusion column (Izon, Medford, MA, USA, Cat # SP5, serial #1000788) according to the manufacturer’s protocol and the 1.5 ml flow through was collected. The sample was then concentrated using an Amicon concentration column (Millipore Sigma, Cat # UFC801024) for 45-60 min at 3,000 × g at RT. Eight replicates of 20 ml media volumes were processed for TSC and EVT EV isolation and then combined into one sample. Only six replicates of ST3D media were combined. This sample was quantified and characterized thrice by Zetaview Nanoparticle Tracking Analysis (NTA; methods described below). The sample was then divided into three aliquots, one each for RNAseq by freezing in Qiazol (Qiagen, Germantown, MD, USA, Cat #217004), mass spectrometry by freezing in PBS (FisherScientific, Cat #BP3991), and Western Blot by freezing in Pierce RIPA buffer (Fisher Scientific, Waltham, MA, USA, Cat #P18990) with 1X HALT (Thermo Fisher, Cat #78440).

### EV sample quantification and characterization

EVs were quantified by Zetaview NTA (Particle Metrix, Meerbusch, Germany) using the following parameters: minimum brightness = 23, maximum size = 800, minimum size = 8, tracelength = 16, nm/class = 4, class/decade = 64, sensitivity = 80.2, frame rate = 30, and shutter = 100. Each sample was analyzed thrice, and the coefficient of variation (CV) was calculated. The CV within each sample set was less than <6%. To determine the percentage of potential Zika virions in the EV samples, the percentage of particles 50-54 nm in size ^36^ was calculated.

To determine if Zika virions were present in the EV preparations or whether the ZIKV envelope (E) protein co-localized with EVs released by infected cells, samples were incubated in AlexaFluor-488 labeled anti-ZIKV E antibody (1:10 dilution) for 2 hrs at RT in the dark prior to quantification with the Zetaview NTA. Samples were diluted at least 1:1000 according to the Zetaview NTA fluorescence protocol. As a positive control, 5.5 ml ZIKV stock was put through the EV isolation protocol, stained with the conjugated E antibody, and evaluated by NTA following the same protocol as described for the EV preparations. The fluorescent settings were minimum brightness = 20, maximum size = 800, minimum size = 5, tracelength = 11, nm/class = 4, class/decade = 64, sensitivity = 80, frame rate = 30, and shutter = 100.

### Transmission electron microscopy

For transmission electron microscopy (TEM) imaging of EVs, 6 EV samples (2 per cell type, 1 ZIKV and 1 control each) were fixed in 2% paraformaldehyde (PFA; Fisher Scientific, Cat #50-980-487) and processed as cited ^59^. Fixed samples were then placed on Formvar-carbon coated electron microscopy grids. After 20 mins the grids were washed with PBS. The grids were then placed in 1% glutaraldehyde for 5 mins and washed in distilled water 8 times. Grids were transferred to a uranyl-oxalate solution (pH 7) for 5 mins and then a methyl cellulose-UA solution for 10 mins on ice. Samples were dried and imaged.

### Detection of ZIKV via immunofluorescence

To evaluate cells exposed to ZIKV by immunocytochemistry, additional cells were infected and cultured as described above. For staining, TSCs were cultured for 60 hrs on col IV coated coverslips (5 μg/ml), rinsed in PBS, and fixed with 4% PFA for 10 mins. Previous work showed that EVTs did not grow well on glass coverslips. Thus, after culture, EVTs were lifted, rinsed in PBS, fixed in 4% PFA, and cytospun at 1000 × g for 1 min onto coverslips. ST3Ds were cultured for 48 hrs, then transferred to wells containing col IV-coated coverslips (5 μg/ml) and allowed to attach for 2 hrs. The cells were rinsed with PBS and fixed in 4% PFA for 10 mins. After fixation, all coverslipped cells were rinsed twice with PBS and stored at 4°C in PBS. For immunostaining, the cells were permeabilized for 10 mins with 0.1% Triton-100 (Millipore Sigma, cat #T-9284) and then blocked for 10 mins with Background Punisher (Biocare Medical, Pacheco, CA, USA, Cat #BP974H). For antibody details please see Supplemental Table 1. Cells were incubated with either primary specific or rabbit IgG isotype control antibody diluted in DaVinci Green Diluent (Biomedical Care, PD900M) for 1 hr at RT, washed 3 times at 5 mins each with 0.1% Tween Tris-buffered saline (TBST), and then exposed to secondary rabbit antibody for 45 mins at RT. Finally, the cells were stained with DAPI for 5 mins, washed in Milli-Q water, and then coverslips were adhered to slides using ProLong Diamond Mountant (Fisher Scientific, Cat # P36961). The following day the sides were imaged using a Nikon confocal microscope and Elements software (Nikon, Tokyo, Japan).

### Lactate Dehydrogenase (LDH) Apoptosis Assay

To determine if ZIKV induced cell death, an LDH (Cytotox96 non-radioactive cytotoxicity assay, Promega, Madison, WI, Cat #G1780) time course was completed on each cell type. Infection of Vero cells was done as a positive control. For this assay, additional cells were inoculated at the same MOI and length of duration as previously stated and shown in Supplemental Figure 1A.

TSCs (100,000 cells/well) were seeded into 24-well col IV coated plates (Corning, Cat #3527). The following day they were exposed to ZIKV or mock infected. EVTs were plated (100,000 cells/well) in collagen IV coated 24-well plates and differentiated in the wells following the previously stated media changes. ST3Ds were differentiated in non-adherent T25 flasks, and after they were exposed to ZIKV (day 3 of differentiation, as previously stated) they were transferred (three-quarters of a T25 flask/well) to non-adherent 24-well plates (Eppendorf, Hamburg, Germany, Cat #0030 722.019). Vero cells were plated in 24-well plates and then exposed to ZIKV at an MOI of 1, 5, or 10 for 1 hr the following day. Cells were plated such that three ZIKV-exposed and three control wells were quantified at each time point.

After the ZIKV exposure (as previously determined for each cell type), cells were rinsed once with PBS and then an LDH assay was performed. For the LDH assay, media were removed, the cells rinsed with PBS, and then 100 μl of 1X lysis solution was added. The cells were incubated in lysis solution at 37°C for 45 mins. After the 45 min incubation, 50 μl was removed and put into a 96-well plate. A volume of 50 μl of the CytoTox 96 reagent was added and the plate incubated for 30 mins at RT in the dark. Finally, 50 μl of Stop Solution was added, and the plate was immediately read at 490 nm. The average among the three wells was calculated and the change in LDH in infected cells compared to controls was determined.

### Plaque assay

To determine the quantity of virus in the conditioned media, plaque assays on Vero cells were conducted as previously reported ^58^. Samples were assessed in duplicate (EVT) or triplicate (TSC and ST3D).

### Hormone quantification

Monkey chorionic gonadotropin (mCG) and progesterone assays were performed as previously reported ^58^. Samples were run in duplicate and unconditioned media were analyzed to determine background mCG and progesterone. The lower limit of detection for the mCG and progesterone assays are 0.1 ng/ml and 10 pg/ml, respectively.

### Total RNA isolation from cells

Total RNA was isolated from cells using an RNeasy kit (Qiagen, Cat #74104) following kit recommendations with modifications. A volume of 700 μl of Qiazol was added to the cell pellet and then frozen at -80°C until RNA extraction. The protocol was followed with the same minor adaptations as with the miRNeasy kit. A 15 min DNAse treatment was performed on the column using RNase-Free DNase (Qiagen, Cat # 79254) prior to washing the column.

### Total RNA isolation from EVs

Each EV sample was diluted in 5 volumes Qiazol and incubated for 5 mins at RT prior to freezing at -80°C until RNA extraction. The miRNeasy serum/plasma kit (Qiagen, Cat #217184) was used to isolate total RNA from EVs following the manufacturer’s protocol with minor adaptations. The Qiazol with Chloroform was overlaid onto phase maker tubes (ThermoFisher, Cat #A33248) and then spun at 4°C at 16,000 × g for 15 mins. In addition, two RPE washes of 500 μl were applied prior to RNA elution. All samples were eluted in 30 μl RNAse free water and the initial elution was placed back on the column and reeluted to increase RNA concentration. Sample concentration and purity was determined with the NanoDrop One (ThermoFisher, Cat # ND-ONE-W).

### Poly(A)RNAseq

RNA integrity was checked with Agilent Technologies 2100 Bioanalyzer. Poly(A) tail-containing mRNAs were purified using oligo-(dT) magnetic beads with two rounds of purification. After purification, poly(A) RNA was fragmented using a divalent cation buffer in elevated temperature. The poly(A) RNA cDNA sequencing library was prepared following Illumina’s TruSeq-stranded-mRNA sample preparation protocol. Quality control analysis and quantification of the sequencing library were performed using Agilent Technologies 2100 Bioanalyzer High Sensitivity DNA Chip. Pair-end sequencing reads of 150 bp reads were generated on an Illumina NovaSeq 6000 sequencing system.

For cell poly(A)seq data, sequencing reads were filtered to remove adaptors and primer sequences and to remove sequences with a quality score lower than 20 ^60^. The cleaned sequencing reads were aligned to the reference genome (GCF_003339765.1_Mmul _10) using the HISAT2 package ^61^. Multiple alignments with a maximum of two mismatches were allowed for each read sequence (up to 20 by default). Transcript abundance estimation and differential expression analysis of aligned reads of individual samples were assembled using StringTie ^62^. Transcriptomes from all samples were then merged to reconstruct a comprehensive transcriptome using a proprietary Perl script designed by LC Sciences. Following transcriptome reconstruction, raw read counts were filtered, normalized, and differential expression determined with DESeq2 ^63^ and edgeR ^64,65^. Genes were considered differentially expressed if they were called by both DESeq2 and edgeR. The glmLRT test was used for edgeR differential expression analysis. EdgeR normalized values were used to produce the heatmaps.

For EV poly(A)seq data, sequences were filtered and adaptors removed with Cutadapt ^60^. Salmon ^66^ was used to obtain transcripts estimates. The mapping rate for the eight sequencable samples was between 3.5-12% for all except one, which had a mapping rate of 86%. Due to the low quality we chose to assess purely based on transcript presence or absence in control/ZIKV EVs. EVs released by control TSCs and ST3Ds (five total samples) were combined and compared to EV data obtained from ZIKV exposed TSCs and ST3Ds (three total samples). Due to limited sample size, differential expression analysis was not performed.

### miRNA-seq

The total RNA quality and quantity was assessed using a Bioanalyzer 2100 (Agilent, CA, USA). Approximately 0.4 - 1 μg of total RNA were used to prepare small RNA cDNA libraries utilizing the specified protocol for the TruSeq Small RNA Sample Prep Kit (Illumina, San Diego, USA). Single-read sequencing of 50 bp was performed on an Illumina Hiseq 2500 at LC Sciences LLC (Houston, Texas, USA). Read trimming and analysis were performed by the University of Wisconsin Biotechnology Center. miRNA abundance estimations were done using the miARma-Seq miRNA-Seq workflow. Accessions for known miRNAs from the Mmul_8.0.1 were obtained from miRbase v22 (October 2018). Data were filtered with the edgeR filterByExpr function and statistical analysis was performed with the edgeRglm function.

### Validation of gene expression by qRT-PCR

A second RNA sample was extracted to generate technical replicates to validate gene expression changes by qRT-PCR that were initially identified by Poly(A)-sequencing analysis. cDNA was synthesized with a SuperScript III First-Strand Synthesis kit (ThermoFisher, Cat # 18080051) using 1 μg RNA per reaction. The manufacturer’s protocol was followed and the Oligo dT primer was used. qRT-PCR was performed with iQ SYBR Green Supermix (Bio Rad, Cat #1708882) following the manufacturer’s recommendation for primer concentration, master mix formulation and primer concentration. Beta-actin mRNA was amplified in parallel to serve as the reference gene and the primer sequences are listed in Supplemental Table 2. cDNA was diluted 1:10 and reactions were run in triplicate. The cycling protocol indicated with the kit was followed. Beta-actin was used to normalize and the 2^-ΔΔCt^ method ^67^ was used to determine changes in transcript abundance.

### Protein quantification and sample preparation

EV and cell protein samples for Western Blots were solubilized in Pierce RIPA buffer with 1X HALT, sonicated for 3-6 sec, spun down at 10,000 × g for 10 mins, and frozen at -80°C. Samples were quantified using the BCA assay as previously described ^12^. For cell samples, 2-4 samples were quantified to estimate the total amount of cell protein in the study.

### Western Blots

EV samples were diluted in 2 or 4X TPA buffer (2X recipe: 20 mM TRIS, 2 mM EDTA, 1 mM Na_3_VO_4_, 2 mM DTT, 2% SDS, 20% glycerol, and a few drops of bromophenol blue) and heated for 5 mins at 70°C. Cluster of differentiation (CD) 9, Heat shock protein 70 (Hsp70), and calnexin were stained for based on the MISEV 2018 guidelines ^68^. To detect ZIKV, the E and nonstructural protein 2B (NS2B) antibodies were used; see Supplemental Table 1 for antibody information.

A total of 3 μg of cell, EV, or an in-house quality control (QC) protein sample was added per well of a 12% polyacrylamide gel along with 8 μl PageRuler Standard (Thermo Fisher, Cat #26616). Samples were prepared such that 2 identical blots were run at the same time. Gels were run in Running buffer (National Diagnostics, Cat # EC870). Protein transfer was done using the GENIE Electrophoretic Transfer (IdeaScientific, Minneapolis, MN, USA) and Transfer buffer (National Diagnostics, Cat # EC880) with 20% methanol (Fisher Scientific, Cat #A412P-4). Protein was transferred onto PVDF paper (Millipore Sigma, Cat # IVPH00010, Billerica, MA, USA) for 90 mins at 0.5 amps with a Power Pac HC power supply (Bio Rad, Cat #1645052). The blots were blocked at 4°C overnight in 5% nonfat dry milk (RPI, Cat # M17200, Mount Prospect, Illinois, USA) in 1X TBST. Blots were incubated in primary antibody (at the concentrations specified in Supplemental Table 1) for 1 hr on a rocker in 5% milk at 37°C. The blots were then washed in 1X TBST three times for 5 mins each at RT and then exposed to either mouse or rabbit secondary antibody for 1 hr on a rocker in 5% milk at 37°C. The blots were washed again in TBST three times for 5 mins each at RT and then exposed to Immobilon Crescendo HRP (Millipore Sigma, Cat # WBLUR0100) for 3 mins at RT and imaged in a Bio-Rad ChemiDoc XRS+ with ImageLab software.

### Mass spectrometry for proteomics

EV preparations were submitted to the University of Wisconsin Mass Spectroscopy/Proteomics Facility through the Biotechnology Center. Isolated EVs in PBS (5 - 10 μg protein) were acidified with trichloroacetic acid (TCA; Fisher Scientific, Cat #A322-100) (10% vol:vol final) to inactivate ZIKV, diluted with water to 200 μl total volume, and then 250 μl of acetone and 50 μl of TCA were added. Samples were sonicated for 2 mins at 6°C in a sonication bath to lyse and release protein content, incubated on ice for 45 mins to facilitate protein denaturation and precipitation, and then spun for 10 mins at 16,000 × g at RT. Protein pellets were washed once each in acetone and methanol and then re-solubilized and denatured in 20 μl of 8 M Urea, 50 mM NH_4_HCO_3_ (pH 8.5), 1mM TrisHCl. Samples were then diluted to 60 μl with 2.5 μl of 25 mM DTT, 5 μl MeOH, and 32.5 μl 25 mM NH_4_HCO_3_ (pH8.5), incubated at 52°C for 15 mins, cooled on ice to RT, and then 3 μl of 55 mM Iodoacetamide (IAA) was added for alkylation and incubated in darkness at RT for 15 mins. Reactions were then quenched by adding 8 μl of 25 mM DTT. Finally, 6 μl of Trypsin/LysC solution [100 ng/μl 1:1 Trypsin (Promega) and LysC (FujiFilm) mix in 25 mM NH_4_HCO_3_] and 23 μl of 25 mM NH_4_HCO_3_ (pH8.5) was added to 100 μl final volume. Digestion was conducted for 2 hrs at 42°C, 3 μl of trypsin/LysC mix was added, and then the digestion proceeded overnight at 37°C. Reaction was terminated by acidification with 2.5% TFA [Trifluoroacetic Acid] to 0.3% final concentration.

NanoLC-MS/MS was completed as previously described ^69^. The Lumos acquired MS/MS data files were searched using Proteome Discoverer (ver. 2.2.0.388) Sequest HT search engine against Uniprot *Rhesus macaque* proteome database (UP000006718, 11/19/2020 download, 44,378 total entries) using previously published settings. Static cysteine carbamidomethylation, and variable methionine oxidation plus asparagine and glutamine deamidation, 2 tryptic miss-cleavages and peptide mass tolerances set at 15 ppm with fragment mass at 0.6 Da were selected. Peptide and protein identifications were accepted under strict 1% FDR cut offs with high confidence XCorr thresholds of 1.9 for z=2 and 2.3 for z=3 and at least 2 PSMs per protein identification. Strict principles of parsimony were applied for protein grouping. Chromatograms were aligned for feature mapping and area-based quantification using unique and razor peptides. Normalization was performed on total peptide amount and scaling on all averages.

### Luminex assay on conditioned media

To quantify secreted immunomodulatory proteins and growth factors, conditioned media were analyzed with the Protcarta 37-plex (ThermoFisher, Cat #: EPX370-40045-901) as previously described ^58^ and a custom Bio-Rad 6-plex based on the Human Inflammation Panel 1 (BioRad, Cat# 171AL001M). For both assays, samples were run in duplicate on a Bioplex 200 instrument (BioRad, Cat #171000201). Data were analyzed with the Bioplex Manager Software. The lower level of quantification for each analyte is listed in the figure legend and is based on the manufacturer’s quantifications.

### DNA preparation

Cell pellets were frozen at -80°C until DNA extraction. DNA was extracted using a Qiagen FlexiGene kit (Qiagen, Cat # 51206) according to the “Isolation of DNA from cultured cells” protocol. Two to four samples were extracted per sample type and DNA was dissolved in 100-500 μl of the kit’s FG3 buffer. If the sample was viscous after the heat incubation, it was sonicated for 3-6 sec on ice. DNA was quantified using a Nanodrop One spectrophotometer.

### Quantification and statistical analysis

Data are represented as the mean ± standard error of the mean (SEM) of four biological replicates. For Zetaview NTA, each sample was processed at least thrice and the CV was calculated. Zetaview NTA data were analyzed in GraphPad Prism using a Kruskal-Wallis test and post-hoc Dunn’s correction. All secretion data were normalized to the total DNA calculated for that sample set. Statistical analysis of secretion data was performed using GraphPad Prism 9.0 (GraphPad Software) by first log transforming the data prior to conducting a one-way ANOVA test with a post-hoc Bonferroni correction applied (p < 0.05).

For the mass spectrometry, Poly(A)seq, and miRNAseq data, significance was determined with an adjusted p-value < 0.05 and 1 < log_2_ fold change < -1. For heatmaps, the rows were organized by hierarchical clustering using agglomerative clustering with Ward’s minimum variance method and the Euclidean distance metric.

## Acknowledgments

Thank you to Dr. Shelby O’Connor and John James Baczenas in the Department of Pathology and Laboratory Medicine at the University of Wisconsin-Madison for sequencing the ZIKV stock used in this study. We would also like to thank Drs. Greg Barrett-Wilt and Greg Sabat at the University of Wisconsin-Madison Biotechnology Center for processing and analyzing the mass spectroscopy samples. We also thank Dr. Mark Berres at the University of Wisconsin-Madison Biotechnology Center for processing and analyzing the miRNAseq data. Thanks to Dr. Randall Massey at the University of Wisconsin-Madison Electron Microscope Core for taking the transmission electron microscope images. We would like to acknowledge Drs. Kristen Bernard and Margaret Petroff for their guidance in viral stock propagation and EV isolation, respectively. This research was funded by NIH grants F31 HD100057 to L.N.B., K99 HD099154 to J.K.S., NIH T32 GM007133 to M.C.M., R21 HD091163 and R01 AI 132519 to T.G.G, and P51 OD011106-54 to the Wisconsin National Primate Research Center. The content is solely the responsibility of the authors and does not necessarily represent the official views of the NIH.

## Data availability

Poly(A)-seq and miRNAseq data have been deposited at GEO (GSE185113 and *awaiting accession ID*, respectively) and are publicly available as of the date of publication. The EV poly(A)-seq data was deposited and is available at GEO (GSE185291). Mass spectroscopy raw data files were uploaded to the Proteome Exchange and are publicly available as of the date of publication (*awaiting accession ID*).

## Supplemental Figures & Tables

**Supplemental Figure 1.**
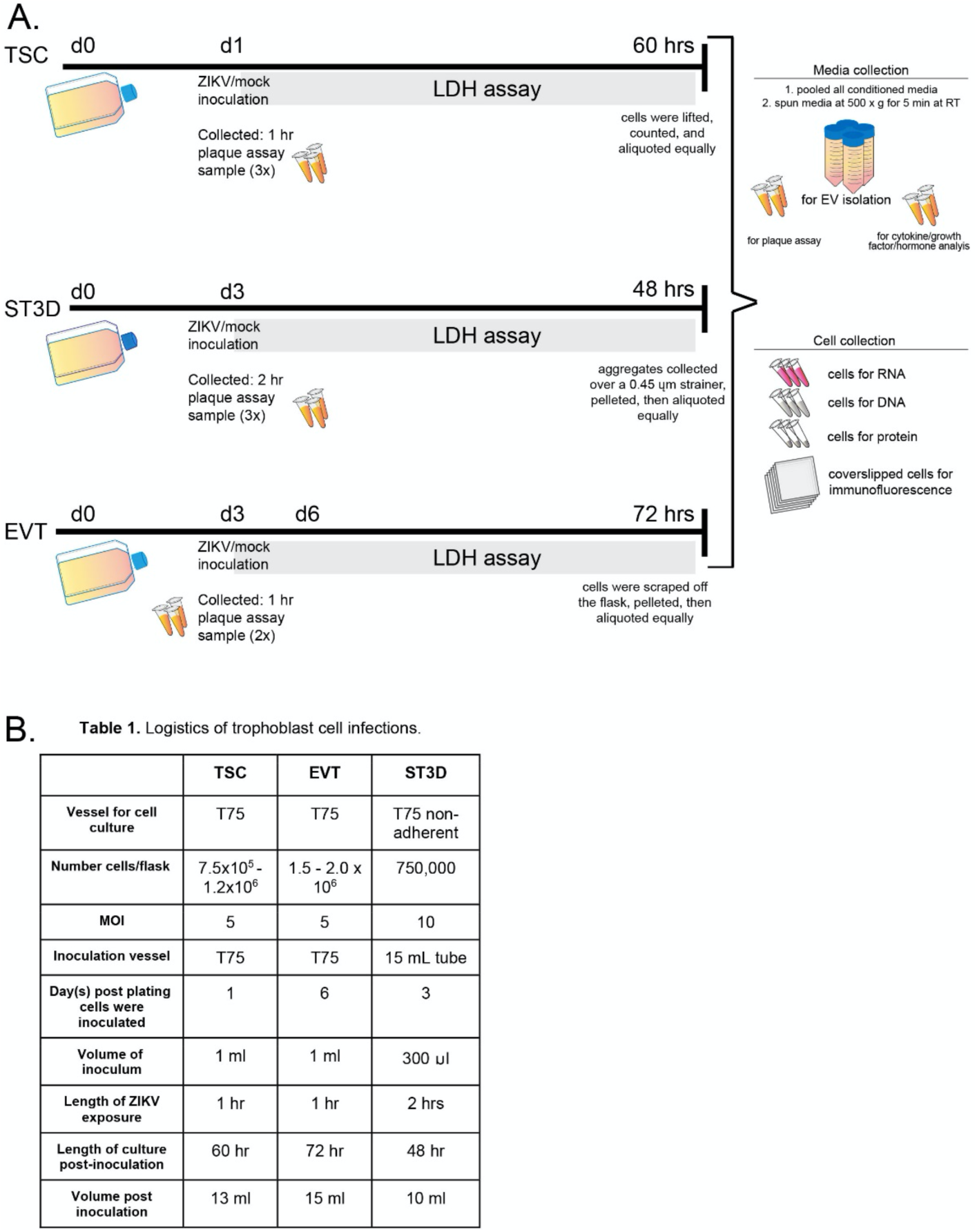
Methods overview. **A)** A timeline depiction of how TSCs, ST3Ds, and EVTs were cultured, when samples were collected, and for what assay samples were collected. **B)** The logistics of trophoblast cell infection including the type of vessel used for cell culture, the number of cells plated, the MOI, inoculation vessel, the day of differentiation cells were inoculated, volume of inoculum, length of ZIKV exposure, length of culture post-inoculation, and media volume post inoculation.

**Supplemental Figure 2.**
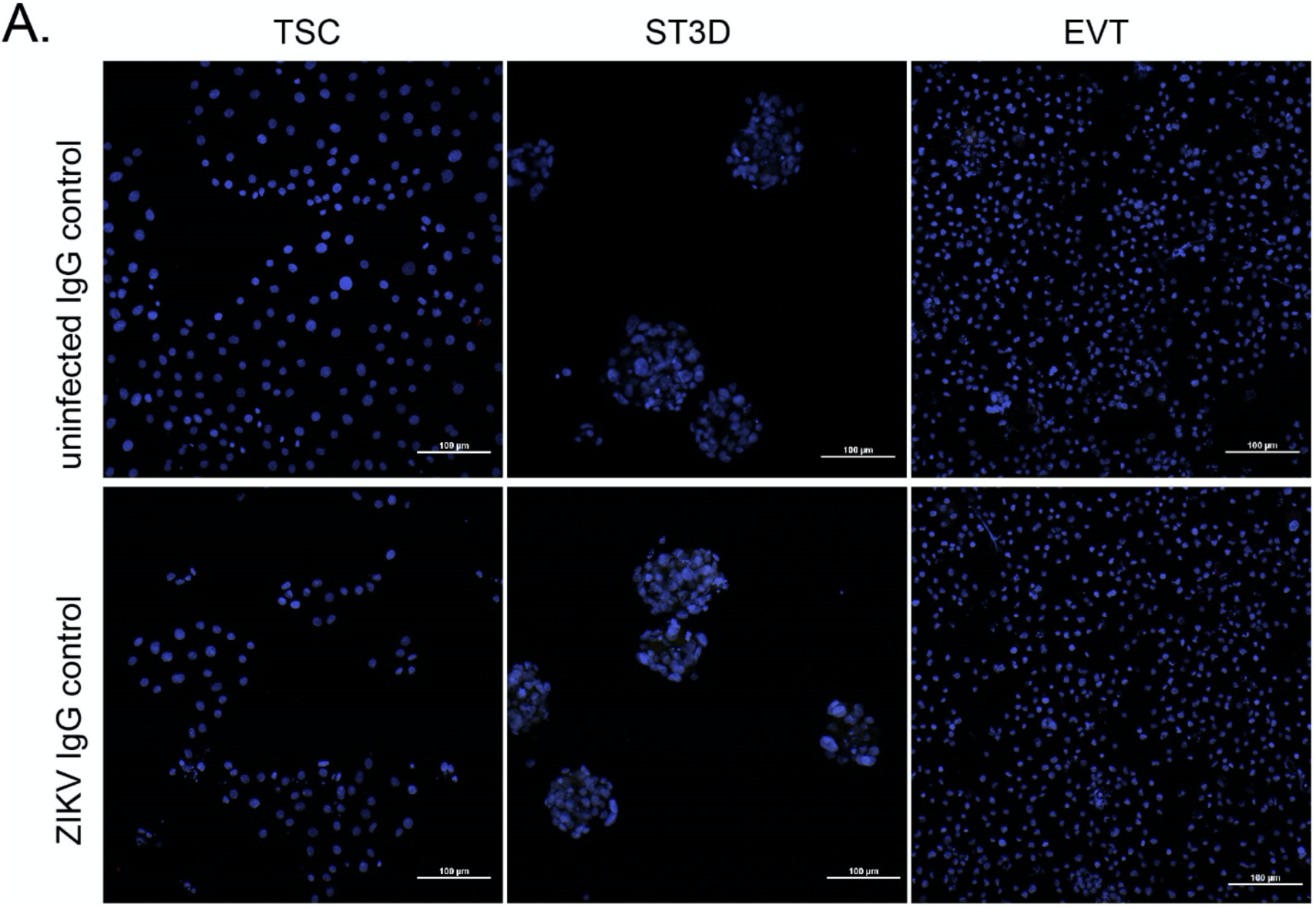
Immunocytochemistry images for rabbit IgG controls and ZIKV uninfected controls. Rabbit IgG controls on TSC, ST3D, and EVT cells.

**Supplemental Figure 3.**
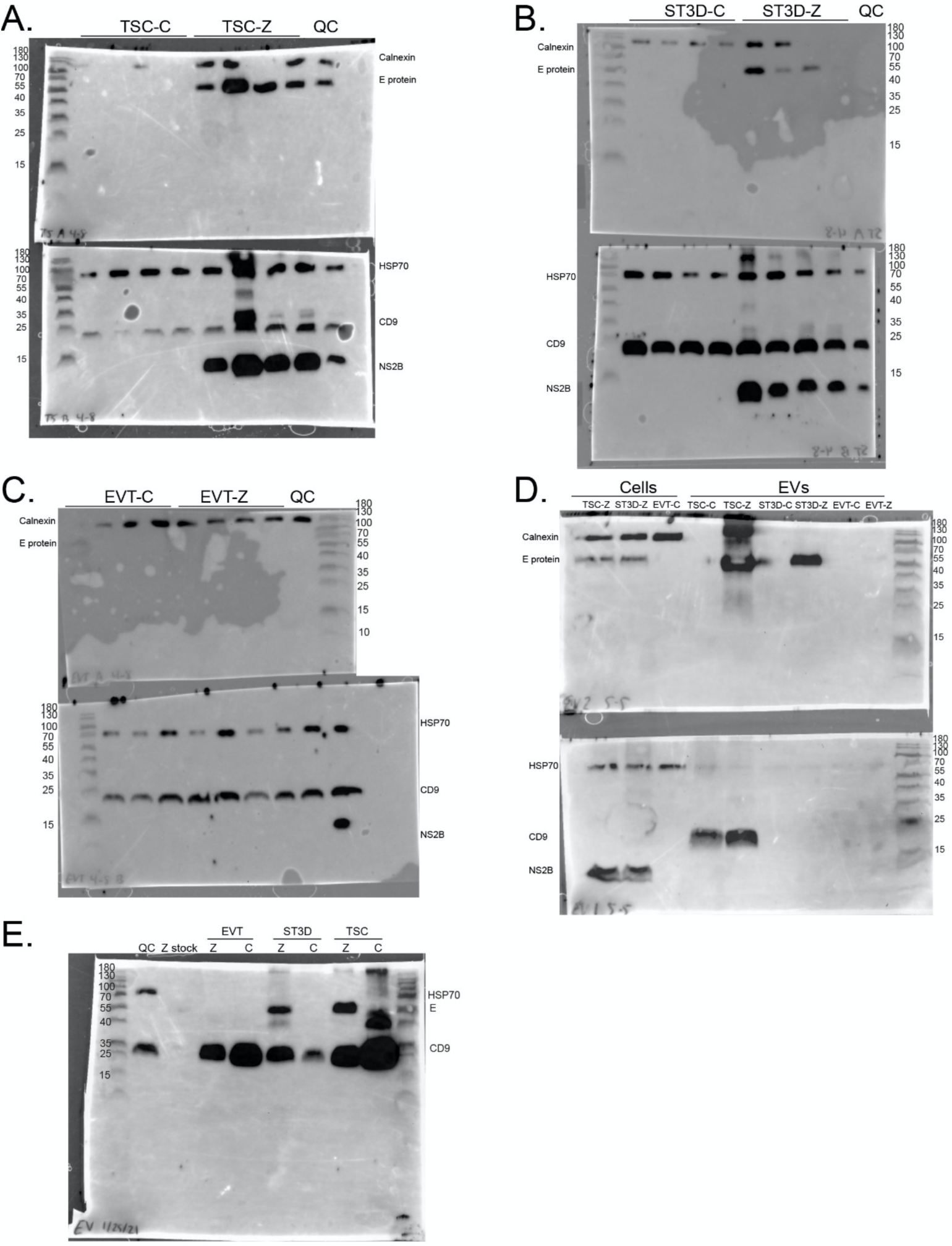
Full Western Blots. Full Western Blots. **A-C)** The full Western Blots of TSCs, ST3Ds, EVTs lysates are shown with the ladder merged with the stained blot. **D)** The Western Blot completed on six EV lysates with three cellular lysates included as a control. **E)** The Western Blot completed on six of the EV lysate samples submitted for mass spectroscopy.

**Supplemental Figure 4.**
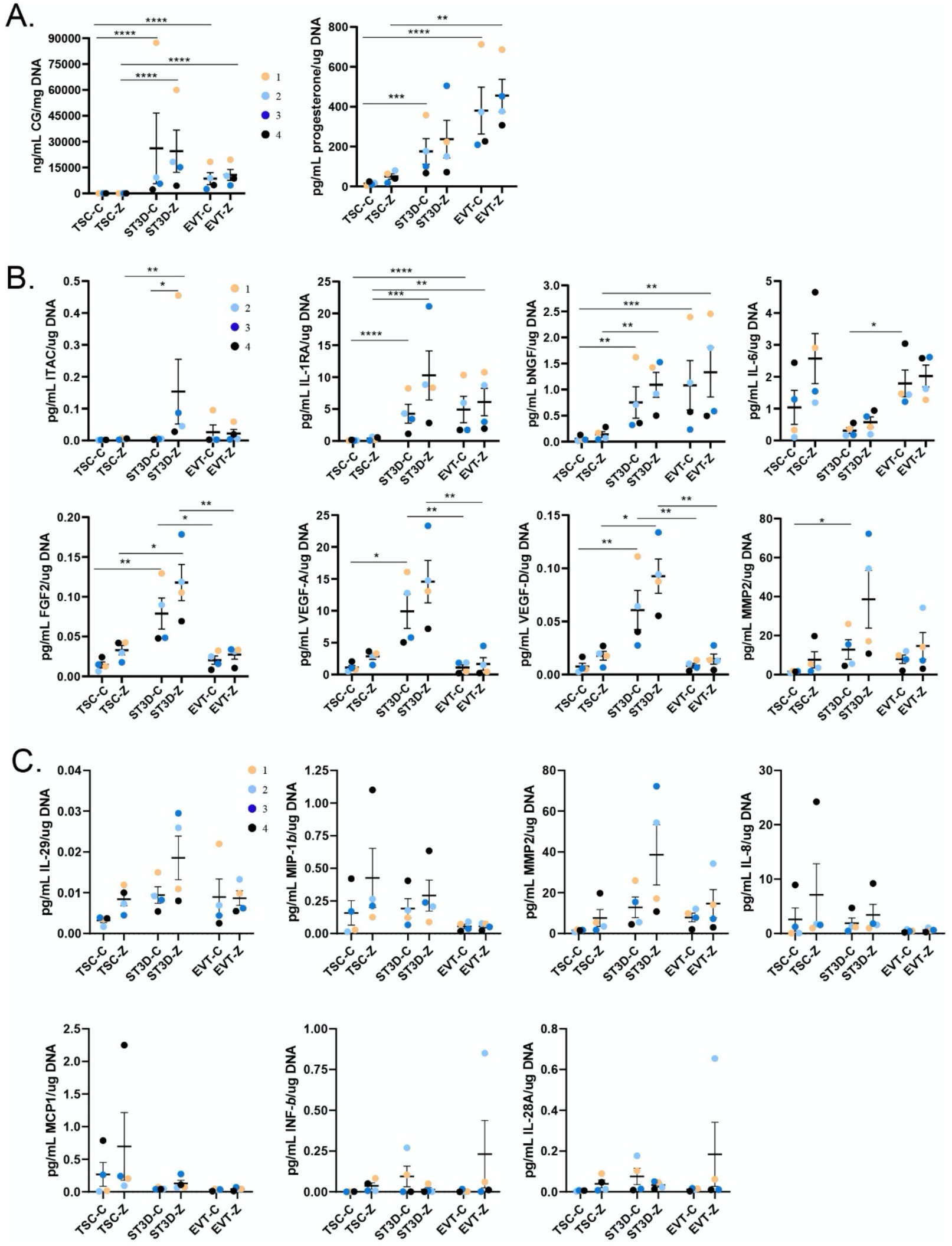
Hormone, cytokine, and growth factor secretion by cell type and infection group. Individual scatter plots of **A)** CG and progesterone and **(B)** cytokines and growth factors are shown. **C)** Luminex assay data on seven analytes (MIP-1b, MCP1, IFN-beta, MMP2, IL-8, IL-28A, and IL-29) which were detected in more than five samples but had no significant changes in expression between ZIKV and control or between cell types. All secretions were normalized to cellular DNA quantity. Significance is depicted on the graphs (**\*** p <0.05; ** p <0.01; *** p <0.001; **** p <0.0001).

**Supplemental Table 1.** Immunocytochemistry, fluorescent conjugates, and western blot antibodies

| Antibody used for | Target | Vendor | Cat number | Clone | Stock Conc | Final conc | Lot number | Isotype |
| --- | --- | --- | --- | --- | --- | --- | --- | --- |
| **IF Primary antibodies** | ZIKV envelope (4G2) | Novus Biological | NBP2-52666 | D1-4G2-4-15 | 1 mg/ml | 1:300 | T1902A20 | Rabbit IgG kappa |
|  | Rabbit IgG | Invitrogen | 31235 |  | 11mg/ml | 1:3300 | VG3028568 |  |
| **IF Secondary antibodies** | donkey anti-rabbit 647 | Jackson ImmunoResearch | 711-605-152 |  | 1mg/800ul | 1:833 | 127614 |  |
|  | donkey anti-rabbit 594 | Jackson ImmunoResearch | 711-585-152 |  | 1mg/800ul | 1:833 | 130064 |  |
| **Other** | Dapi diacetate | Invitrogen | D3571 |  | 10 mg/ml | 1 ug/ml | 1890543 |  |
| **Zetaview antibody** | ZIKV envelope | Novus Biological | NBP2-52709AFF488 | 4G2-4-15 | 0.68 mg/ml | 1:6.67 | T1908B05-020321-AF488 |  |
| **Western blot primary antibodies** | CD9 | Invitrogen | MA1-80307 | MM2/57 | 1 mg/ml | 1:2000 | UL2889567 & UC2744222 | Mouse |
|  | ZIKV E | GeneTex | 133314 |  | 0.45mg/ml | 1:900 | 43306 | Rabbit |
|  | Hsp70 | GeneTex | GTX111088 |  | 0.73 mg/ml | 1:730 | 43082 | Rabbit |
|  | NS2B | GeneTex | GTX133308 |  | 1.15 mg/ml | 1:1150 | 42564 | Rabbit |
|  | Calnexin-CT | StressMarq Biosciences | SPC-108B |  | 1 mg/ml | 1:1000 | PC185088 | Rabbit |
| **Western blot secondary antibodies** | anti-mouse | Jackson ImmunoResearch | 115-035-146 |  | ----- | 1:20,000 | 151651 |  |
|  | anti-rabbit | Jackson ImmunoResearch | 111-035-144 |  | ----- | 1:20,000 | 151080 |  |

**Supplemental Table 2.**
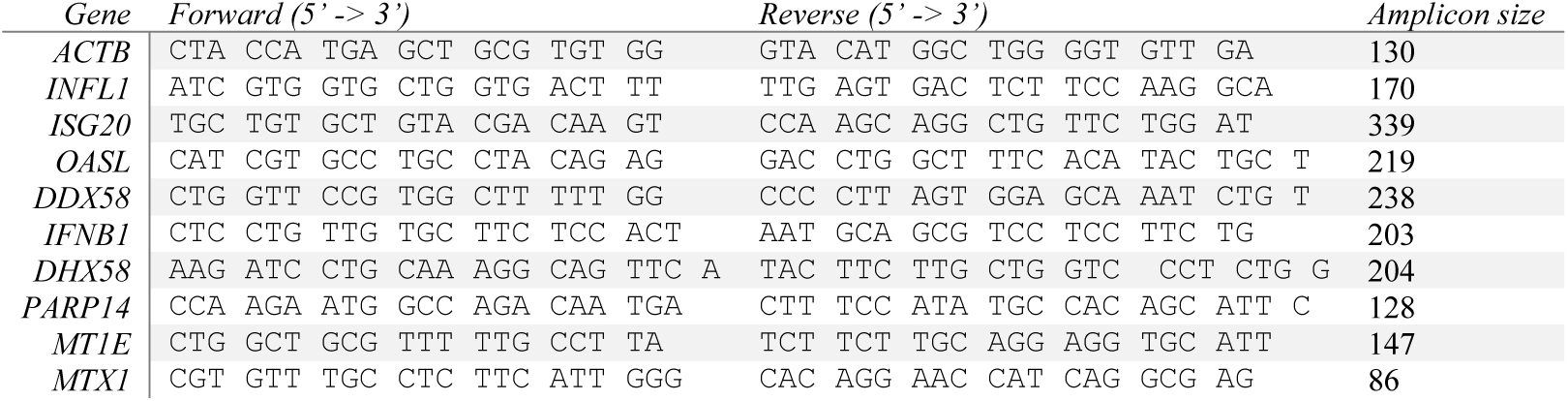
Primer sequences for qRT-PCR.

**Supplemental Table 3.**
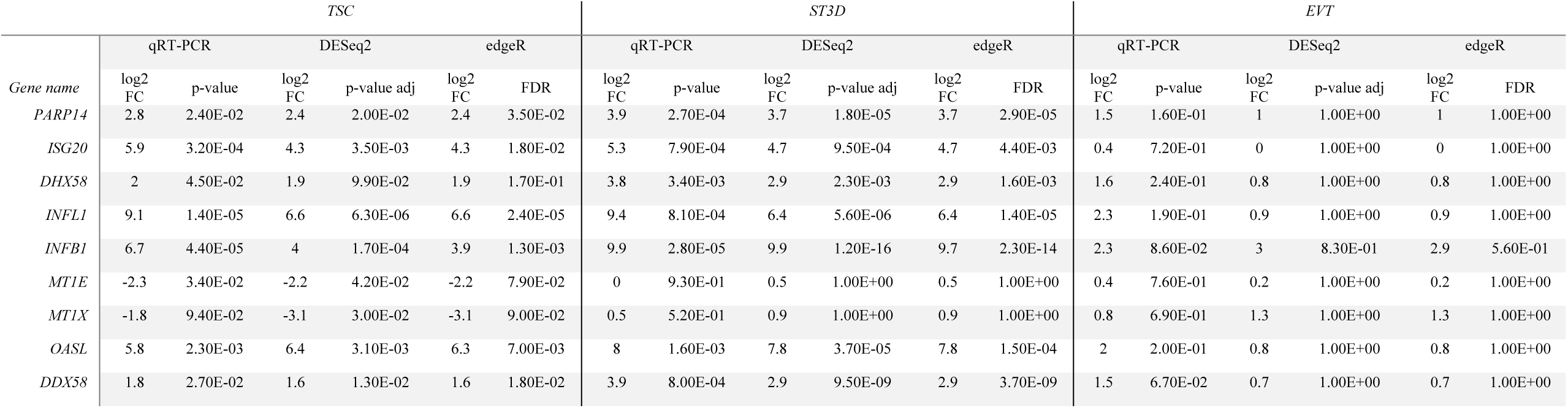
qRT-PCR, DESeq2, and edgeR log fold change (log2FC) and significance (p-value/adjusted p-value) comparisons

